# Genome-wide analysis of lncRNAs points to their roles in the modulation of developmental regulator expression during plant male germline development

**DOI:** 10.1101/2022.08.03.502631

**Authors:** Neeta Lohani, Agnieszka A. Golicz, Annapurna D. Allu, Prem L. Bhalla, Mohan B. Singh

## Abstract

LncRNAs can function in regulating of gene expression, but their roles as essential regulators of developmental processes and organismal phenotypes remain largely unclear. Especially the roles of lncRNAs in plants are largely unexplored. However, it has been proposed that plant lncRNAs act as regulators of protein-coding genes during development and that the similar roles of animal and plant lncRNAs result from convergent evolution. Since pollen development follows an established program with well-defined and characterized stages, we have used it as a model for studying plant lncRNAs and their roles in reproductive development. We investigated of lncRNA expression and function during pollen formation in field mustard (*Brassica rapa*). Reference-based transcriptome assembly performed to update the existing genome annotation identified novel expressed protein-coding genes and long non-coding RNAs (lncRNAs), including 4,347 long intergenic non-coding RNAs (lincRNAs, 1058 expressed) and 2,045 lncRNAs overlapping protein-coding genes on the opposite strand (lncNATs, 780 expressed). The analysis of expression profiles reveals that lncRNAs are significant and stage-specific contributors to the gene expression profile of developing pollen. Gene co-expression networks accompanied by genome location analysis identified 38 cis-acting lincRNA, 31 cis-acting lncNAT, 7 trans-acting lincRNA and 14 trans-acting lncNAT to be substantially co-expressed with target protein-coding genes involved in biological processes regulating pollen development and male lineage specification. These findings provide a foundation for future research aiming at developing strategies to employ lncRNAs as regulatory tools for gene expression control during reproductive development.

## Introduction

Transcriptomic analyses have identified gene expression profiles during pollen development of male reproductive tissues from multiple plant species, including Arabidopsis and rice [1-4]. As pollen matures, the transcriptome becomes less complicated, with an increase in the number of stage-preferential transcripts. Pollen is produced in anthers, the male reproductive organs of flowers, and its formation is the conclusion of a highly specialised and strictly regulated developmental cycle in angiosperms [5, 6]. Pollen/microspore mother cells (also known as meiocytes) undergo meiosis to form tetrads of haploid microspores, which then divide mitotically and differentiate, giving rise to the sperm cell-carrying mature pollen. Mature pollen, which represents highly reduced haploid male gametophyte, contains plant male germ cells and is indispensable to sexual reproduction. The stages of pollen development are well defined with stage-specific markers making it an ideal system for studying plant developmental processes [7, 8].

Recently, long non-coding RNAs (lncRNAs) have emerged as important, stage-specific regulators of developmental processes in animals and plants [9, 10]. No lncRNA conservation between plants and animals has been reported. Still it is postulated that lncRNAs can be universal regulators of developmental processes and that their similar functions and mechanisms of action could be a result of convergent evolution [10]. lncRNAs are RNA molecules with more than 200 base pairs in length, lack open reading frames more than 100 amino acids long, and have no protein-coding potential. The discretionary length limit defining lncRNAs distinguishes them from small non-coding RNAs, including microRNAs (miRNAs), small nucleolar RNAs (snoRNAs), and small interfering RNAs (siRNAs). LncRNAs, which are primarily intergenic ncRNAs (lincRNAs), intronic ncRNAs (incRNAs), or natural antisense transcripts (NATs), often show polyadenylation and tend to have highly tissue-specific expression [11]. They act as decoys, molecular scaffolds, or target mimics of miRNAs and siRNA precursors to influence gene expression [12, 13]. When acting as decoys, certain lncRNAs can bind with transcription factors, thereby precluding their interaction with DNA to promote the expression of target genes, while as molecular scaffolds, they can bind with DNA or protein-recruiting regulatory components to specific gene loci [12-14].

Several reports have highlighted the critical role of lncRNAs in plant biological processes such as stress response, development regulation, and nutrient procurement by regulating modification of histones, transcription, alternative splicing, chromatin remodeling, or target mimicry [11, 15-20]. In Arabidopsis, during cold exposure, an antisense transcript - *COOLAIR* (*Cold Induced Long Antisense Intragenic RNA*) - and an intronic lncRNA - *COLDAIR* (*COLD ASSISTED INTRONIC NONCODING RNA*) - restrict the transcriptional activation of the floral repressor *FLOWERING LOCUS C* (*FLC*) via histone modification and thereby promote flowering [21-23]. Similarly, another cold-induced natural antisense lncRNA, *MAS* (*MAF4 antisense RNA*), is reported to direct the activation of *MADS AFFECTING FLOWERING4* (*MAF4*) via histone modification resulting in the suppression of early flowering in Arabidopsis [24]. In rice, the silencing of an antisense lncRNA – *LRK Antisense Intergenic RNA* (*LAIR*) – results in reduced plant growth along with reduced expression of *LEUCINE-RICH REPEAT SERINE/THREONINE-PROTEIN KINASE* (*LRK*) gene cluster [25]. Lines overexpressing *LAIR*, on the other hand, show a significant increase in overall grain yield and increased expression of some members of the *LRK* gene cluster. It was reported that in rice, *LAIR* could variably activate the promoters of the *LRKs* gene by binding to histone modification enriched in the *LRK1* gene area [25].

In plants, lncRNAs have also been linked to male reproductive development. In rice, under transcription of long-day conditions, long-day–specific male-fertility–associated RNA, *LDMAR* is required for photoperiod-sensitive male sterility (PSMS) activation and proper pollen formation [26]. In young panicles of rice, overexpression of *LDMAR* impairs fertility under long-day conditions. In maize, high expression of lncRNA *Zm401* was observed in developing male gametophytes and mature pollen, and it was identified as the primary regulator of genes essential for pollen formation, such as *ZmC5, ZmMADS2*, and *MZm3–3* [27]. Downregulation of *Zm401* leads to aberrant tapetum and microspore development, resulting in the production of sterile pollen. Furthermore, in Chinese cabbage (*Brassica campestris* L.), a novel pollen-specific lncRNA-*BcMF11* was identified to regulate male reproductive development [28, 29]. The silencing of BcMF11 resulted in delayed tapetum degradation, abnormal microspore development and pollen abortion. These findings demonstrated that lncRNAs are essential for regulating pollen formation.

Here, we performed a genome-wide identification of lncRNAs during five stages (pollen mother cell, tetrad, microspore, bicellular pollen and mature pollen) of pollen development in field mustard (*Brassica rapa*) using strand-specific RNA sequencing (ssRNA60 Seq). lncRNAs exhibit stage-specific expression suggesting potential roles at well-defined developmental points. Next, we analysed the genomic location of lncRNAs and predicted *cis* and *trans*-acting lncRNAs and their potential target protein-coding genes. Differential expression and functional enrichment analysis highlighted the complex transcriptional reprogramming involved in the transition of diploid pollen/microspore mother cells into haploid trinucleate pollen. We further performed a weighted gene co-expression network analysis (WGCNA) coupled with gene expression correlation to identify lncRNA-mRNA pairs with a potential role in regulating pollen development progression. Collectively, our findings shed light on the roles of lncRNAs during pollen development and expand our knowledge of the molecular mechanisms underlying male reproductive development.

## Results

### Identification and characterization of lncRNAs in *B. rapa* expressed during pollen development

Strand-specific RNASeq sequencing reads corresponding to five stages of pollen development (pollen mother cell – ‘PMC’, tetrad – ‘TET’, microspore – ‘MIC’, binucleate pollen – ‘BIN’ and trinucleate pollen – ‘POL’), were used to track changes in gene expression during male gametophyte development in *B. rapa* (Figure 1A). Both poly(A) capture and ribosomal RNA (rRNA) depletion libraries were prepared. The reads were aligned to the *Brassica rapa* genome with a mapping rate for poly(A) capture libraries between 78.18% and 90.85 % (mean: 87.34%) and for the rRNA depletion libraries between 71.02% and 83.22% (mean: 76.74%; Table S1A). Because pollen development requires the participation of highly specialized tissues and cell types, some of the genes involved may not be found in the existing annotation. A reference-based [30] transcriptome assembly was performed to update the existing genome annotation (using an in-house pipeline, Figure S1, [31]), identify novel expressed protein-coding genes and long non-coding RNAs (lncRNAs), including long intergenic non-coding RNAs (referred to as ‘lincRNAs’ hereafter) and lncRNAs overlapping protein-coding genes on the opposite strand (referred to as ‘lncNATs’ hereafter). In total, 49,577 protein-coding genes, 4,347 lincRNAs and 2,045 lncNATs were identified. Comparison of the poly(A) capture and rRNA depletion libraries (TPM < 0.1 for all the poly(A) libraries and TPM >0.1 in at least one rRNA depletion library) suggests that 1.3%, 4.3%, and 1.7% of loci produce non-polyadenylated transcripts for coding, lincRNA and lncNAT genes respectively. Principal component analysis (PCA) revealed high relatedness between the replicates of each sample (Figure S2A). Further, the Pearson correlation between the three biological replicates ranged from 0.911 to 0.989 (median: 0.968). The correlation between the coding genes, lincRNAs and lncNATs was also significant across the five pollen developmental stages (Figure S2B). We have tested the concordance between expression observed in this dataset and the previously reported expression patterns of known male development markers in *A. thaliana*. All the markers, other than AtMGH3 and AtGEX2, for which no confident orthologues were identified, had expected expression patterns (Figure 1B and S2C).

**Figure 1.**
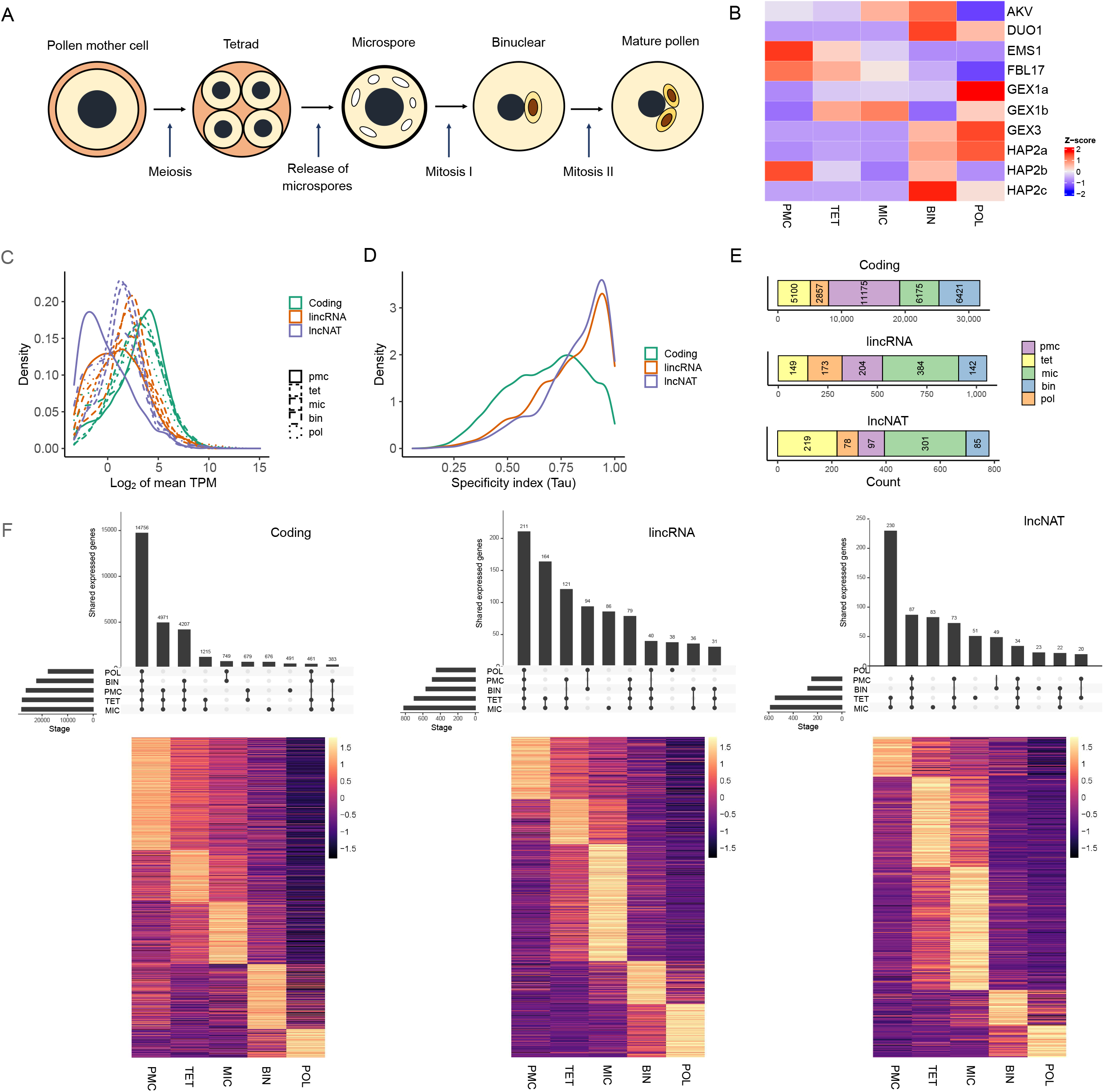
Male gametophyte development and properties of the lncNATs and lincRNAs discovered **(A)** The five stages of pollen development, **(B)** Heatmap of *B. rapa* homologs of known pollen development marker genes, **(C)** Distribution of expression values for coding genes and lncRNAs, **(D)** Expression specificity index for protein-coding genes and lncRNAs, **(E)** Summary of the number of coding and lncRNA genes showing peak expression at a given stage, and **(F)** Heatmaps and Upset plots presenting overall expression patterns of coding genes lncNATs and lincRNAs across the five stages. **PMC**: pollen/ microspore mother cell, **TET**: tetrads, **MIC**: microspores to polarised microspores, **BIN**: early to late bi-nucleate pollen, and **POL**: tri-nucleate pollen.

Lastly, based on data mean-variance trend analysis, genes with low expression were filtered, and a CPM cut-off (>1.0 CPM in at least three samples) was imposed, identifying 31,729 coding genes, 1,052 lincRNA and 780 lncNAT loci available for the analysis. A comparison of the protein-coding and lncRNA loci confirms that the latter have lower expression levels and more stage-specific expression (Figure 1C and Figure 1D), with different expression profiles of coding genes and lncRNAs. Among the samples used in this study, the highest number of protein-coding genes (35.22%), had peak expression in PMC and lincRNAs (36.50%), and lncNATs (38.59%) had peak expression in MIC (Figure 1E and 1F). It is important to note that the peak expression stage has been defined as the stage with maximum gene expression measured by TPM (transcripts per million). Therefore, the Peak expression stage is the stage where transcript abundance is the highest relative to the abundance of other transcripts at that stage.

The lncRNAs were shorter than coding genes with ∼80% and 40% lncRNAs with one transcript and only one exon, respectively (Figure 2A, 2B and 2C). Compared to lincRNAs, lncNATs had slightly higher proportion of lncNATs had one transcript (lincRNAs: 78.61%, lncNATs: 82.18%) and multiple exon (lincRNAs:52.96%, lncNATs: 55.28%). A/U content of the lincRNAs and lncNATs (particularly the lincRNAs) was also higher than the protein-coding sequences (Figure 2D). Among the lncRNAs with assigned chromosome locations (Figure 1E), most expressed lncRNA loci (164 lincRNAs and 107 lncNATs) were mapped to chromosome A09, and the least was found to be present on chromosome A10 (48 lincRNAs and 65 lncNATs). The majority of the expressed mRNA loci were located on chromosome A03 (4618) and the least on chromosome A04 (2212) (Figure 1E).

**Figure 2.**
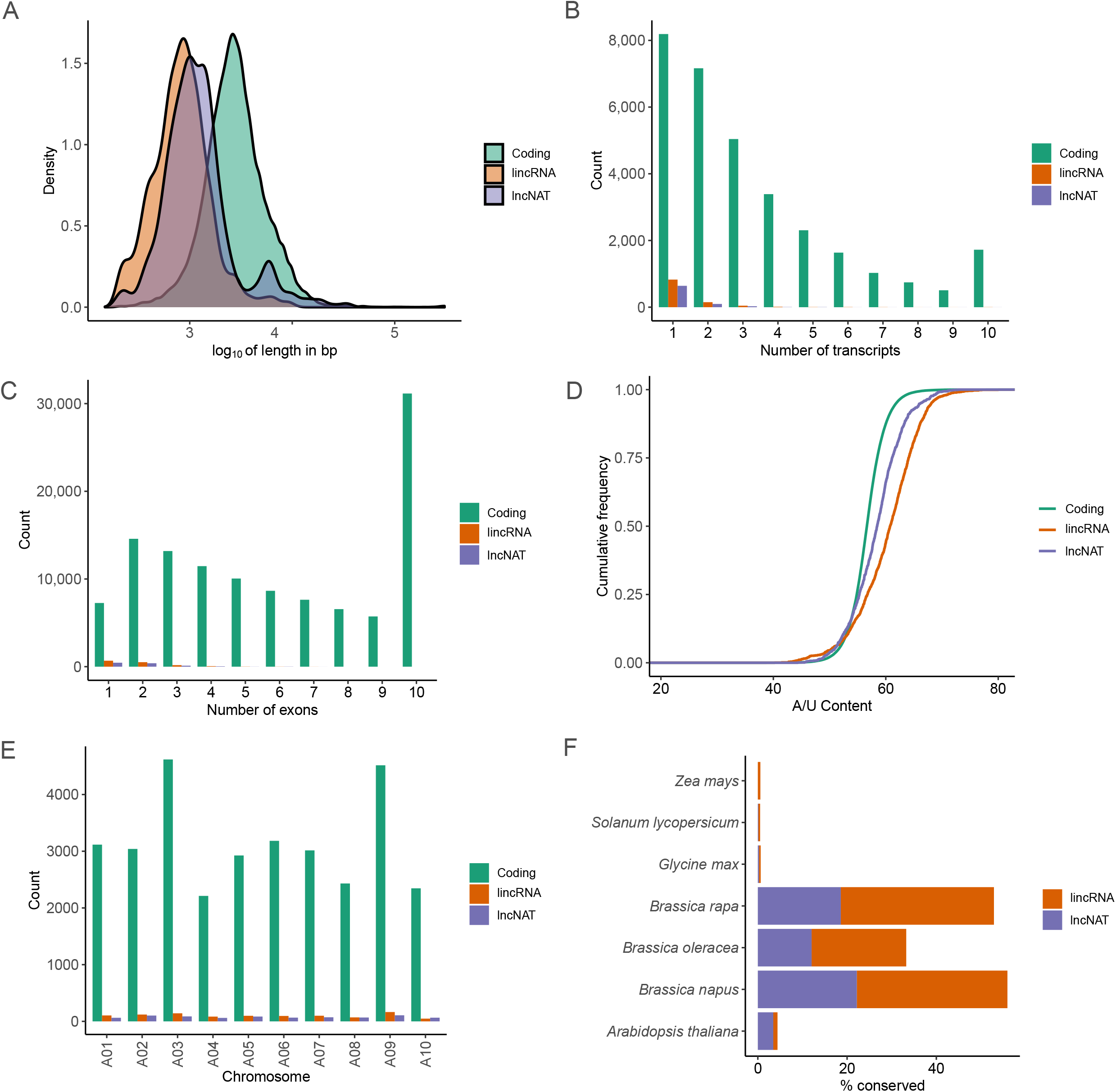
**(A)** Distribution of transcript length of coding genes and lncRNAs, **(B)** Number of transcripts per gene of coding genes and lncRNAs, **(C)** Number of exons per transcript of coding genes and lncRNAs, **(D)** Comparison of A/U content of coding transcripts and lncRNAs **(E)** Chromosome distribution of coding genes and lncRNAs, and (F) Conservation of identified lincRNAs and lncNATs in *B. rapa* and other plant species.

### Conservation analysis of lncRNAs expressed during pollen development in *B. rapa* against other plant lncRNAs

Sequence conservation analysis of the lncRNA sequences identified in this study was performed by BLAST analysis against the reported lncRNAs (retrieved from CANATAdb [32]) from *Arabidopsis thaliana* (4,373), *B. napus* (12,010), *B. rapa* (8,501), *B. oleracea* (7,338), *Glycine max* (3,096), *Solanum lycopersicum* (4,716), and *Zea mays* (10,761). The results indicate that *B. rapa* lncRNAs identified during pollen developmental stages are not well conserved in *A. thaliana, G. max, S. lycopersicum* and *Z. mays*. However, they still show significant conservation between previously published lncRNAs from other *Brassica* species (Figure 1F). However, the level of conservation between the lncRNAs identified in this study and lncRNAs in *B. napus, B. rapa* and *B. oleracea* collectively is still low (12.05% to 33.75%), which highlights the identification of novel male reproductive development specific lncRNAs in *B. rapa* in this study.

### Prediction of *cis-* and *trans-*acting lncRNAs

In the next step, the *cis* and *trans* interactions of the lncRNAs with the expressed protein-coding genes were predicted. The relative location of lncRNA to their neighbouring protein coding gene has been shown to be associated with the effect the lncRNA has on protein-coding gene expression [33]. The *cis*-acting lncRNAs are divided into several classes based on the direction (sense or antisense), type of interactions (intergenic or genic) and relative location (upstream or downstream) with respect to the interacting protein-coding gene [34]. Figure 3A and 3B summarises the *cis* lincRNAs and lncNATs present on A01 to A10 chromosomes, respectively. In this analysis, the lincRNAs are identified as intergenic, and their distribution between sense and antisense is roughly equal (Figure 3A, Table S2). A slightly higher number of lincRNAs are located upstream (2540) of the protein-coding genes compared to the lincRNAs located downstream (1947). lncNATs are identified as antisense and genic, the majority of which are located in exons of protein-coding genes (Figure 3B, Table S3).

**Figure 3.**
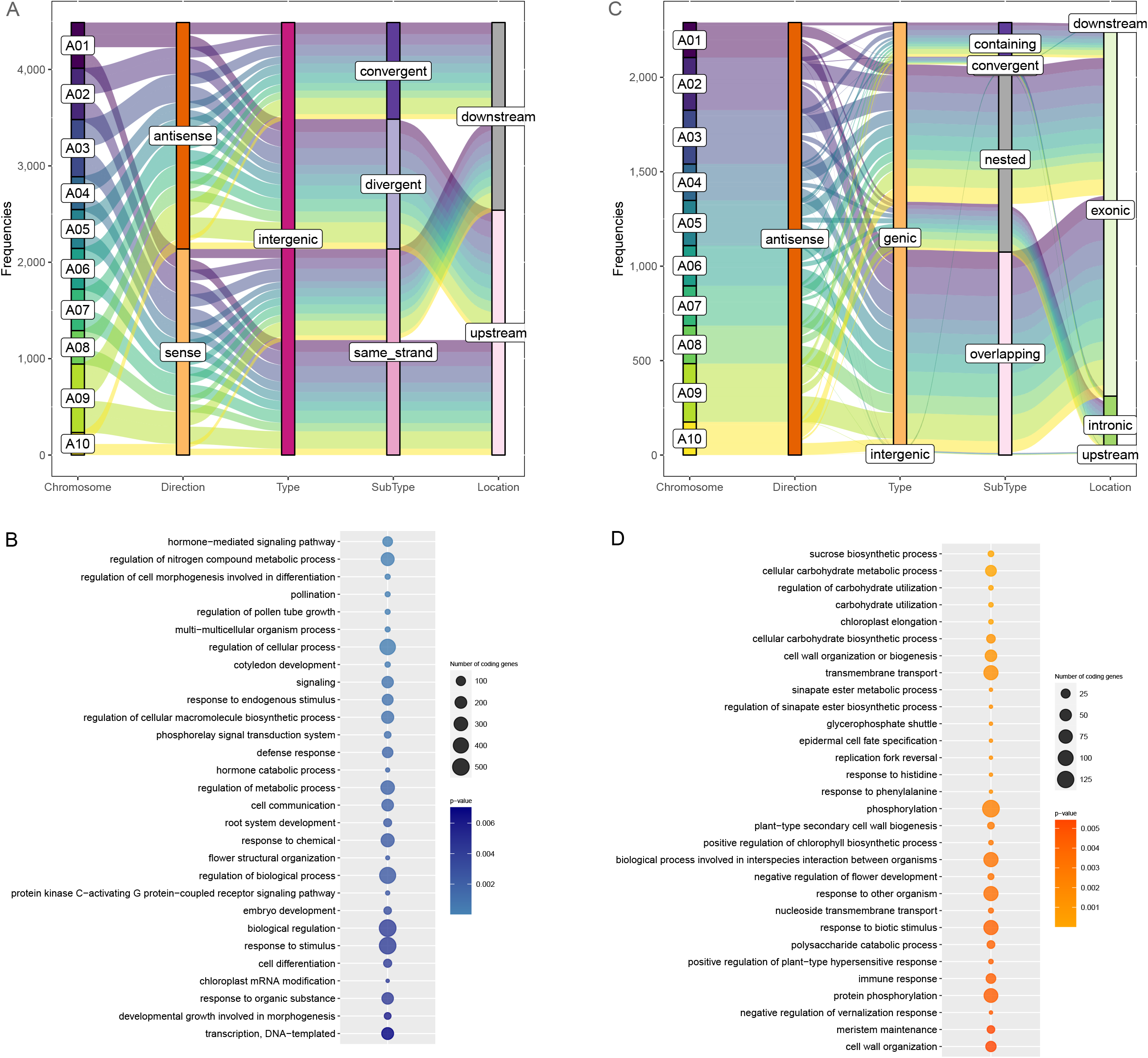
**(A)** lincRNA *cis* interactions classification per chromosome in *B. rapa*, **(B)** Top significant non-redundant GO terms associated with expressed protein-coding genes identified as partners of *cis*-acting lincRNAs, **(C)** lncNAT *cis* interactions classification per chromosome in *B. rapa*, and **(D)** Top significant non-redundant GO terms associated with expressed protein-coding genes identified as partners of *cis*-acting lncNATs.

Further, these *cis*-acting lncRNA-protein coding genes neighbouring pairs were filtered out to select the pairs in which both lncRNA and protein-coding genes were identified as expressed in the samples. The GO enrichment analysis of the protein-coding genes identified as partners of the *cis*-acting lincRNAs and lncNATs is provided in Figures 3C and 3D, respectively. lincRNAs neighbouring proteins coding genes were associated with biological process categories such as “hormone-mediated signalling pathway”, “regulation of pollen tube development”, “cell communication”, “regulation of cell morphogenesis involved in differentiation” and “transcription, DNA-templated”, (Figure 3C, Table S4). The protein-coding genes neighbouring *cis*-acting lncNATs were involved in “carbohydrate utilization”, “transmembrane transport”, “replication fork reversal”, “phosphorylation”, and “stamen filament development” among other biological processes (Figure 3D, Table S5).

The prediction of *trans* regulation of protein-coding genes by lncRNAs depends on the formation of complementary hybrids and the associated interaction energy between the lncRNA and the associated protein-coding genes. Interactions in the scaffold were discarded since the scaffold is unplaced, and one cannot determine the *bona fide* of the *trans* interactions. Initially, the maximum threshold of interaction energy was set at -20 joule to retain significant interactions, and 103,545 interactions were identified for lincRNA transcripts. For lncNAT transcripts, 82,606 total *trans* interactions were identified. However, a number of significant *trans* interactions in the order of hundreds of thousands are unlikely. The distribution of the energy of interactions (Figure 4A), shows that most of these interactions have low energy (below -100 joule). Thus, setting a more stringent arbitrary threshold of -100 joule (red vertical line in Figure 4A) brings down the number of *trans* interactions to 1,418 for lincRNAs and 1,061 for lncNATs. The 1,418 identified *trans* interactions involved 548 lincRNAs (Table S6), out of which ∼43% significantly interacted with only one protein-coding gene, whereas nine lincRNAs interacted with ≥10 protein-coding genes. *LINC_BRAPST00049411* interacted with the maximum number of protein-coding genes (39) in a *trans* manner. In contrast, 1061 *trans* interactions involved 575 lncNATs (Table S7), with only 65% lncNATs interacting with one protein-coding gene, and nine lncNATs interacted with ≥10 protein-coding genes. Among the lncNATs, *NAT_BRAPST00007879* interacted with 28 protein-coding genes. Further, these lncRNA-protein coding genes *trans* interacting pairs were filtered out to select the pairs in which both lncRNA and protein-coding genes were identified as expressed in the samples.

**Figure 4.**
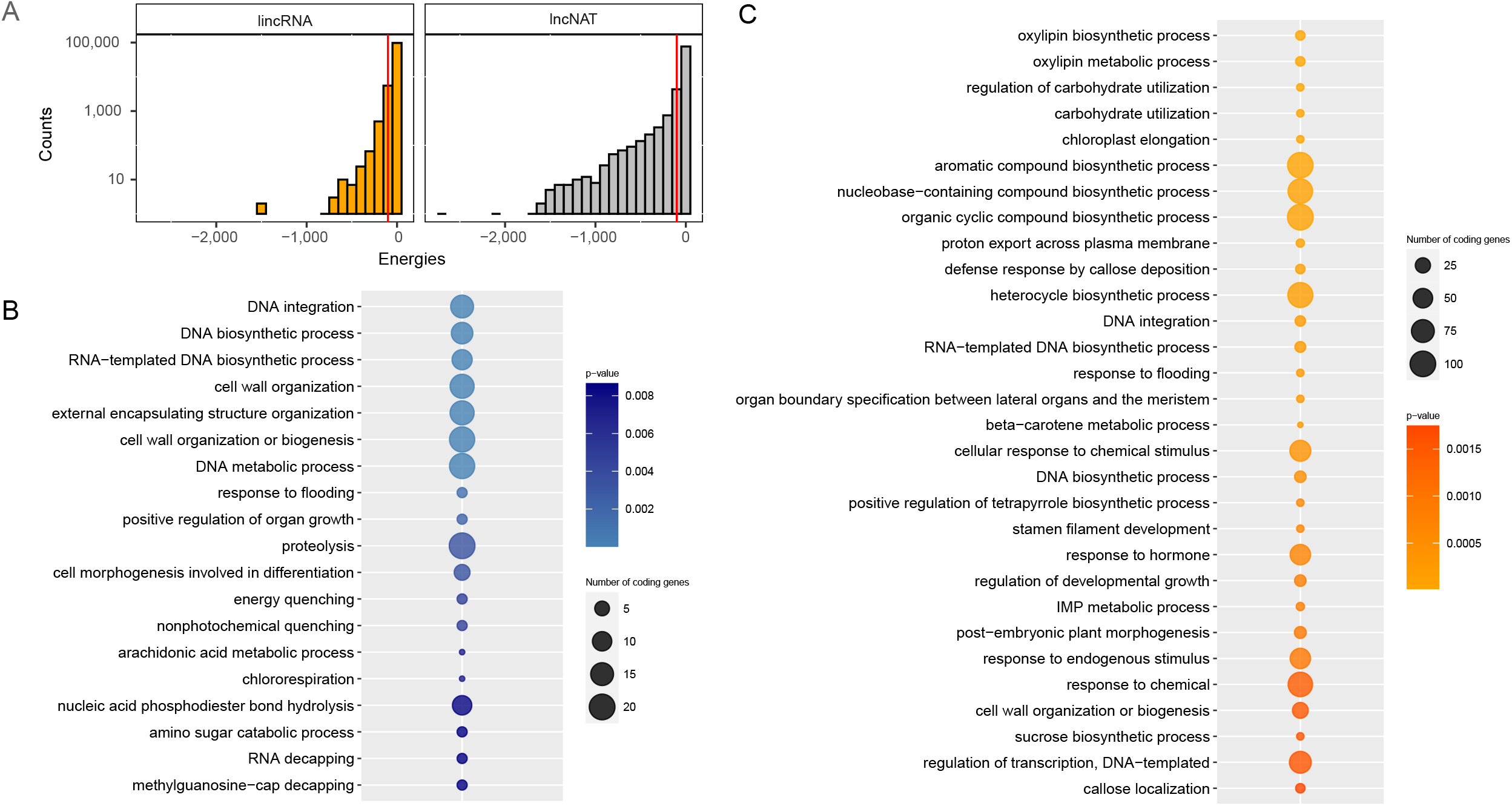
**(A)** Comparison of *trans* interactions free energy distribution for lincRNAs (LINC) and lncNATs (NAT), **(B)** Top significant non-redundant GO terms associated with expressed protein-coding genes identified as partners of *trans*-acting lincRNAs, and **(C)** Top significant non-redundant GO terms associated with expressed protein-coding genes identified as partners of *trans*-acting lncNATs.

Functional enrichment of protein-coding genes identified as potentially regulated by lincRNAs in a *trans* manner revealed their association with “DNA integration”, “cell wall organization”, “proteolysis”, “cell morphogenesis involved in differentiation”, and “regulation of cell growth” among other biological process categories (Figure 4B). Furthermore, *trans*-acting lncNATs potentially regulated protein-coding genes involved in biological processes such as “oxylipin biosynthetic process”, “carbohydrate utilization”, “DNA integration”, “stamen filament development”, and “response to hormone” (Figure 4C).

### lncRNA as potential miRNAs targets and precursors

microRNAs (miRNAs) play an important role in regulating gene expression by influencing mRNA degradation and translational repression [35]. lncRNAs, like mRNA, can be miRNA targets and operate as miRNA decoys, suppressing the interaction between miRNAs and their target genes [13]. Out of the 1,052 lincRNAs, only 22 were predicted as potential targets of 18 miRNAs, and 21 out of 780 lncNATs were predicted to be targeted by 36 miRNAs (Table S8). The majority of the identified *B. rapa* lncRNAs targeted by miRNAs were potentially regulated by cleavage, and very few lncRNA were inhibited at the translational level. The low number of lncRNAs detected as miRNA targets in this analysis is probably due to the lack of male reproductive tissue-specific miRNAs available in published miRNAs.

Some lncRNAs are also considered small RNA (miRNA and siRNA) precursors. We compared the lncRNA sequences to the miRbase collection and found that only 0.95% of lincRNA and 0.90% of lncNATs are potential small RNA precursors (have 100% similarity to known mature miRNAs). Furthermore, to identify high confidence targets of the miRNAs for which lncRNAs served as precursors, psRNATarget with a stringent expectation cut-off of 0 was employed (Table S8). Three lncRNA-miRNA-mRNA modules were identified (Figure 5A). For one of the modules, the expression profile of the lncRNA was antagonistic to the protein-coding gene expression profile. *LINC_BRAPST00004757* acts as a precursor of *bra-miR162-3p*, which then targets the expression of *BRAPST00013543*. Functional annotation identified *BRAPST00013543* as a gene encoding thymidine kinase that salvages DNA precursors. The pyrimidine salvage pathway is crucial for genome replication and maintaining of its integrity. *BRAPST00013543* showed the highest expression in the PMC stage, and its expression gradually decreased, whereas *LINC_BRAPST00004757* expression increased as pollen development progressed (Figure 5A). Thus, it can be postulated that *LINC_BRAPST00004757* regulates the expression of *BRAPST00013543* during male gametophyte development in *B. rapa*.

**Figure 5.**
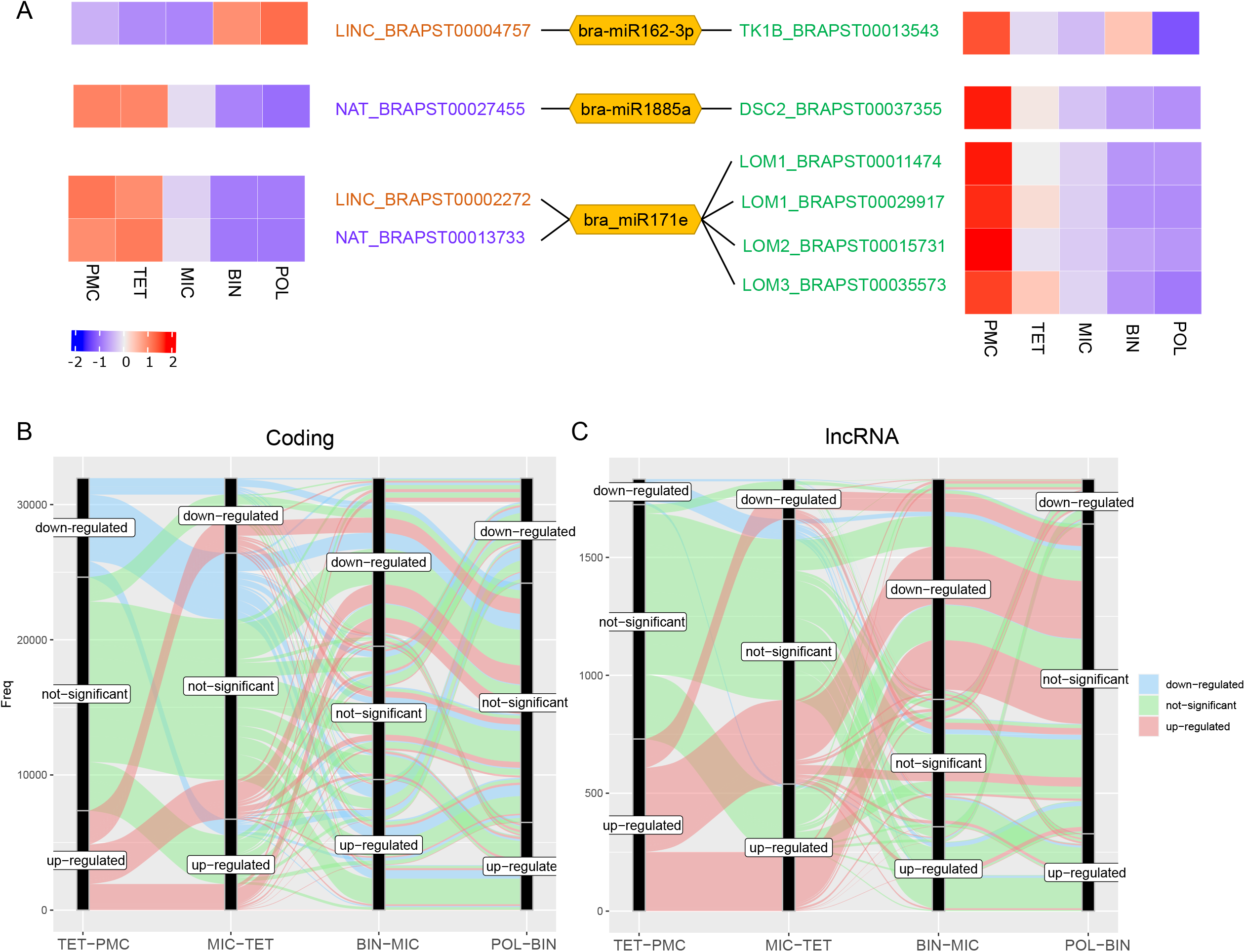
**(A)** Three identified lncRNA-miRNA-mRNA modules, where the lncRNA acts as a precursor of the miRNA, and the mRNA or the protein-coding gene is the direct target of miRNA. Heatmaps represent the expression profiles of lncRNA and protein-coding genes during pollen development, **(B)** Differential regulation of protein-coding genes during pollen development as depicted by an alluvial plot, and **(C)** Differential regulation of lncRNAs during pollen development as depicted by an alluvial plot.

### Differential transcriptional reprogramming during pollen development

LncRNAs identified in the datasets used in this study had lower expression levels than protein-coding genes. LncRNAs with low abundance might get filtered out while performing differential expression analysis (Assefa et al., 2018). *limma* R package employed in this study to perform differential expression analysis runs a moderated t-test after an empirical Bayes correction (Ritchie et al., 2015), a generic and suitable for the differential expression of processed lncRNA expression data. In the RNA-Seq libraries, 49,577 protein-coding genes, 4,347 lincRNAs and 2,045 lncNATs were identified. A CPM cut-off of 1 in at least three samples was used to identify 31,729 coding genes, 1,052 lincRNAs and 780 lncNATs expressed during pollen development. To investigate the regulation of protein-coding genes and lncRNA during pollen development, we performed differential expression analysis (log^2^fc cut-off = 0.585, adjusted p-value cut-off <0.01) across four contrasts (TET-PMC, MIC-TET, BIN-MIC, and POL-BIN) by comparing each pollen developmental stage with the previous one (Table S9, Figure S3A). In total, 92.58%, 89.73% and 93.21% of the expressed protein-coding genes, lincRNAs and lncNATs, respectively, were differentially regulated across the four contrasts. When the uninucleate microspore transitions into a binucleate microspore, a significantly higher percentage of genes and lncRNAs were differentially regulated with a higher proportion of downregulated genes (Figure 5B and 5C, Figure S3A). These observations align with the reported findings that during male germline development, a decreasing trend in transcriptome size and complexity throughout microsporogenesis and microgametogenesis is observed in flowering plants [2, 36]., Only 19 and 84 genes were commonly up- or down-regulated among the protein-coding genes across all developmental stage contrasts (Figure S3B). Interestingly, no common lncRNAs were identified to be differentially regulated across all four contrasts, further highlighting their stage-specific regulation (Figure S3B).

Gene ontology (GO) analysis of the differentially expressed protein-coding genes was performed to unravel their role during different pollen developmental stages (Figure S4). When pollen/microspore mother cell transitions into tetrads, protein-coding genes associated with biological process categories such as “mRNA processing”, “transcription by RNA polymerase II”, “proteolysis”, “gene expression” and “histone modification” were upregulated. Further, as the pollen development progresses, biological process categories including “ribosome biogenesis”, “ncRNA processing”, “translation”, and “gene expression” were upregulated. As highlighted earlier, gene expression significantly downregulates as the uninucleate microspore transitions into binucleate pollen, contributing to the decreasing complexity of transcription in binucleate pollen. This was further supported by the downregulation of biological process categories involved in “regulation of transcription, DNA-templated” and “regulation of gene expression”. In contrast, the upregulated genes were associated with “cellular localization” and “protein transport” among others. During the final trinucleate pollen stage, the genes involved in transcription and protein synthesis were generally expressed at lower levels, as indicated by the downregulation of GO terms “ribosome biogenesis”, “translation”, and “mRNA processing”. Furthermore, protein-coding genes involved in processes including “localization”, “regulation of pollen tube growth”, “ion transmembrane transport”, and “nucleotide-sugar metabolic process” were upregulated in trinucleate pollen. The functional annotation of differentially expressed genes highlighted the stage-specific differential regulation of an array of biological processes during the progression of male gametophyte development.

### lncRNA-mRNA co-expression analysis

To predict the regulatory roles of lncRNA during pollen development, the co-expression networks between the protein-coding genes and lncRNAs (expressed genes, >1 CPM in at least three samples) were identified using the WGCNA tool. The tool identified 24 modules in the dataset (Figure S5). For further analysis, the top three modules associated with the five pollen developmental stages were identified (Table S10). A different number of lncRNAs were present in the selected modules (Table S10). Based on the *cis* and *trans* regulation of protein-coding genes by lincRNA and lncNATs, we next investigated the hub genes in the selected modules and identified lncRNA-protein coding genes interactions and grouped them as *cis*-lincRNA-protein coding gene, *cis*-lncNAT-protein-coding gene, *trans*-lincRNA-protein coding gene, and *trans*-lncNAT-protein coding gene co-expressed pairs. We also performed correlation analysis to supplement the WGCNA analysis and further filtered out lncRNA-coding gene pairs with a Pearson correlation coefficient of less than 0.8 or more than -0.8. Additionally, only those pairs were selected in which the protein-coding gene was differentially expressed and had >65% similarity with its homolog in *A. thaliana*. In total, 54 *cis*-lincRNA-protein-coding genes, 58 *cis*-lncNAT-protein-coding gene, 8 *trans*-lincRNA-protein-coding genes and 18 *trans*-lncNAT-protein coding gene interacting co-expressed pairs were identified (Table S11).

We further searched for an association between those genes and lncRNAs expressed during male gametophyte development based on functional annotation of genes. We collected genes annotated with the GO biological process terms associated with male gametophyte and pollen development. We also collected genes predicted to be transcription factors or homologous to pollen-specific genes in *A. thaliana*. In total, we found 38 *cis*-lincRNA-protein-coding genes, 31 *cis*-lncNAT-protein-coding genes, 7 *trans*-lincRNA-protein coding genes and 14 *trans*-lncNAT-protein-coding gene pairs of interest (Figures 6 and 7). Several key pollen developmental regulators were found among the genes identified, including genes involved in “regulation of cell cycle”, “microtubule-based movement”, “pollen development”, “pollen tube growth”, “cell wall organization”, and “transmembrane transport” along with genes showing pollen-specific expression (Figure 6 and 7). Analysis of the function of protein-coding genes in the identified lncRNA-protein coding gene pairs revealed genes involved in transcription regulation, such as transcription factors belonging to WRKY, bHLH and NAC TF families (Figure 7). The proximity of lncRNAs and our previous results suggesting a possible regulatory relationship between lncRNAs, and their co-expressed protein coding gene partners suggest that the expression of these developmental regulators and transcription factors could be affected by lncRNAs and that they present targets for future investigation.

**Figure 6.**
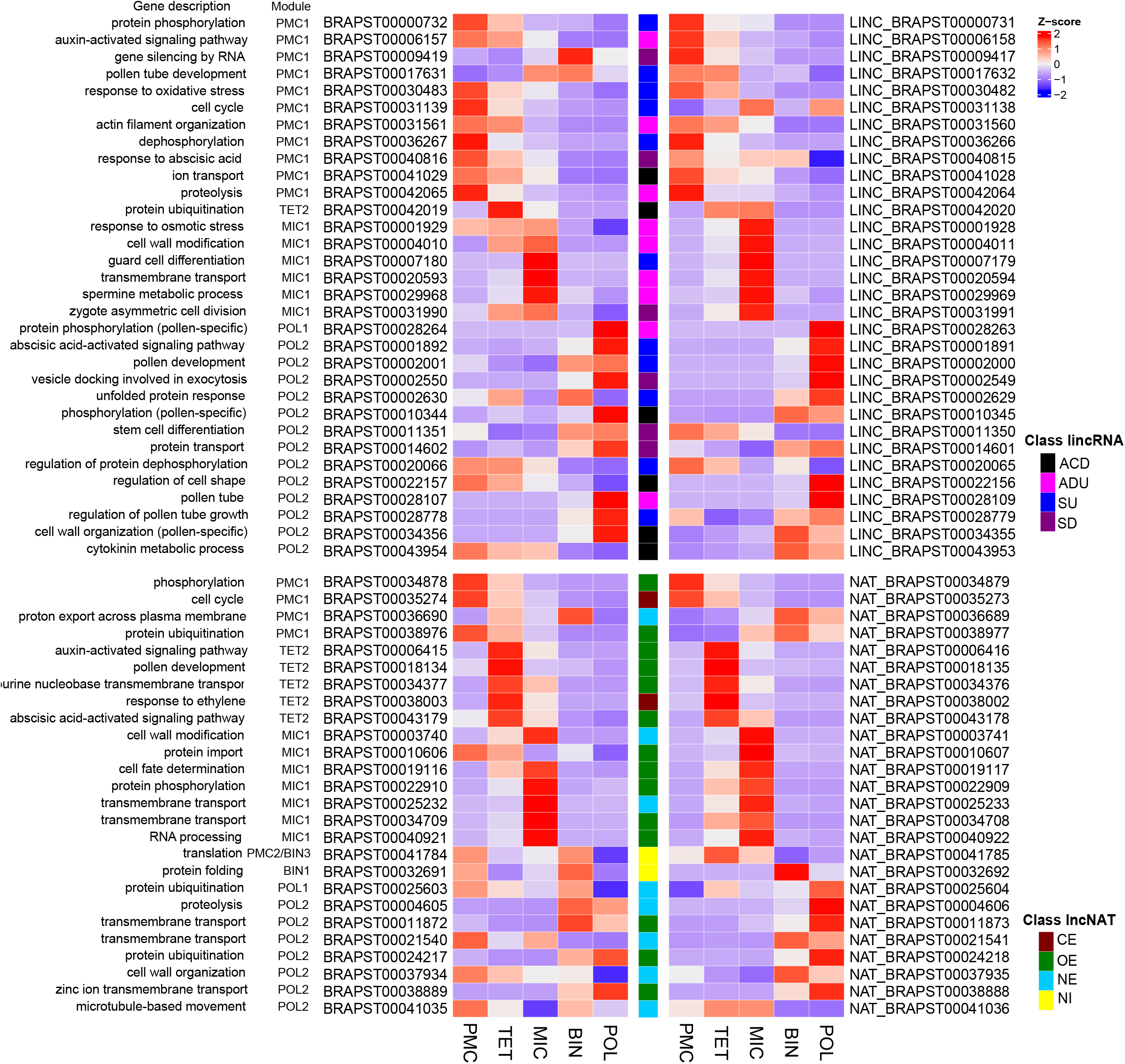
Heatmap representing the *cis*-lincRNA-protein coding gene and *cis*-lncNAT-protein coding gene pairs in the top modules of pollen development stages as identified by the co-expression analysis. The GO term description of the protein-coding gene highlights the involvement of the co-expressed pairs in biological processes critical to pollen development. The class of *cis* interaction between the lncRNAs, lncNATs and their respective co-expressed protein-coding gene is also illustrated. *cis* Classification of lincRNAs-ACD: Antisense, Convergent, Downstream; ADU: Antisense, Divergent, Upstream; SU: Same strand, Upstream; SD: Same strand, Downstream. *cis* Classification of lncNATs-CE: Contained, Exonic; OE: Overlapping, Exonic; NE: Nested, Exonic; NI: Nested, Intronic.

**Figure 7.**
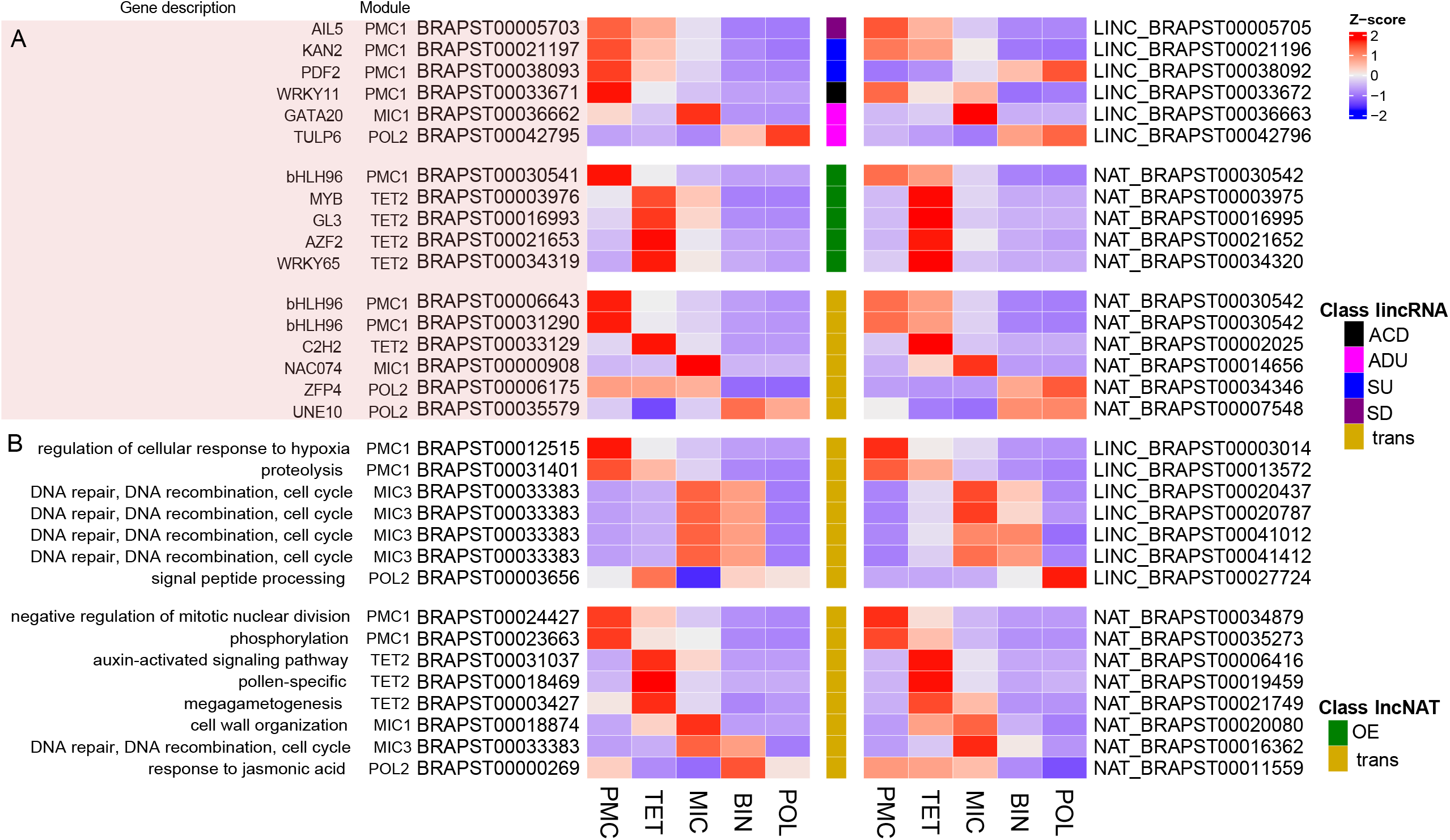
**(A)** Heatmap representing the *cis*- and *trans-* lincRNA-protein coding gene lncNAT-protein coding gene pairs in the top modules of pollen development stages as identified by the co-expression analysis. The protein-coding genes in these pairs were identified as transcription factors (light red box), **(B)** Heatmap representing the *cis*-lincRNA-protein coding gene and *cis*-lncNAT-protein coding gene pairs in the top modules of pollen development stages as identified by the co-expression analysis. The GO term description of the protein coding gene highlights the involvement of the co-expressed pairs in biological processes critical to pollen development. The class of *cis* interaction between the lncRNAs, lncNATs and their respective co-expressed protein coding gene is also illustrated. *cis* Classification of lincRNAs-ACD: Antisense, Convergent, Downstream; ADU: Antisense, Divergent, Upstream; SU: Same strand, Upstream; SD: Same strand, Downstream. *cis* Classification of lncNATs-CE: Contained, Exonic; OE: Overlapping, Exonic; NE: Nested, Exonic; NI: Nested, Intronic.

## Discussion

Long non-coding RNAs (lncRNAs) play diverse roles in regulating of biological processes [9, 10, 37, 38]. Recent research has shown an increasing number of lncRNAs with significant tissue-specific expression patterns, implying that they play regulatory roles in developmental processes [39-41]. Plant reproductive development is an essential feature of crop breeding, and identification of lncRNAs that influence reproductive development is becoming increasingly important. The progression of pollen development from diploid pollen mother cells to haploid microspores to highly specialized haploid trinucleate pollen provides the opportunity for dissecting molecular program controlling male lineage development and identification of transcriptional network in each stage associated with cell identity, cell cycle, cell fate determination as well as male gametophyte specification processes. Here, we identified sets of lncRNAs and protein-coding genes expressed during pollen development (Figure 1A) in field mustard (*B. rapa*). As an important vegetable crop, *B. rapa* is an attractive option for use as a reference species in *Brassica* genome investigations [30]. We predicted regulatory roles of lncRNAs based on genome location and co-expression analysis. Our results help resolve the distinctive identity of pollen developmental stages and provide a rich data source for further describing the mechanisms underlying male lineage development.

We identified 6,392 lncRNAs (4,347 lincRNAs and 2,045 lncNATs) during pollen development in *B. rapa* (Figure 1E). A study in *B. rapa* conducted a time series of RNA-seq experiments at five developmental stages during pollen development and three different time points after pollination and identified 12,051 putative lncRNAs [42]. This difference in results can be attributed using different cell types for sequencing. Huang et al. [42] used whole buds representing the five pollen developmental stages for sequencing, whereas we isolated pollen mother cells, tetrads, microspores, binuclear pollen and trinucleate pollen for sequencing. Another reason can be the use of a different variety of *B. rapa* and the employment of different bioinformatic tools for identificating and discovering of lncRNAs.

The lncRNAs (lncNATs and lincRNAs) identified in *B. rapa* during pollen development (Figure 2A-C) were shorter and had fewer isoforms and exons per transcript than protein-coding genes; these properties were consistent with previous reports of genome-wide lncRNA discovery [16, 41, 43, 44]. About half of the identified lncRNAs reported in Arabidopsis and rice have one transcript and include only a single exon [16, 44]. A higher A/U content is another feature of lncRNAs observed in *B. rapa* lncRNAs (Figure 1D). The A/U composition may reflect the underlying sequence. However, it is attractive to speculate that it may be a feature of lncRNAs that facilitates recognition by RNA-binding proteins. The flexible nature of lncRNAs in interacting with other transcripts is potentially indicated by a high A/U content [45], as transcripts rich in A/U content are less stable [46].

lncRNAs are reported to be low in abundance and show tissue-specific expression in plants and animals [10]. In this study also, *B. rapa* lncRNAs showed lower expression in comparison to protein-coding genes (Figure 1B). The expression patterns of lncRNAs (Figure 1C-F) likely reflect their stage-specific roles during pollen development and are consistent with previous observations of high lncRNA transcription in buds containing the microspore stage [42]. LncRNAs showed peak expression at the microspore stage, whereas a higher fraction of protein-coding genes had peak expression at pollen/microspore mother cells (Figure 1E). Few lncRNAs were also expressed exclusively at a single pollen developmental stage (Figure 1F). Furthermore, differential expression analysis revealed no common *B. rapa* lncRNA to be differentially regulated between all pollen developmental stage contrasts. Different expression windows for lncRNAs suggest they are part of a coordinated expression program rather than a result of non-specific pervasive transcription. These results indicate that lncRNA expression profiles may precisely determine the male lineage specifications. Further investigation of these stage-specific lncRNAs may lead to a better understanding of the molecular control of pollen development.

Across plant species, lncRNAs are reported to show low conservation and are mostly species-specific [47, 48]. Because lncRNAs are highly evolved, there is less sequence conservation across plant and animal taxa, resulting in fewer phylogenetic connections [47]. The present study observed a high divergence between the identified lncRNAs in *B. napus* and reported lncRNAs in *A. thaliana* (Figure 2F). Comparatively higher conservation of lncRNAs were observed between the lncRNA sequences identified in *B. rapa* and previously reported lncRNAs *Brassica* species (*B. rapa, B. oleracea* and *B. napus*) [32]. These lncRNA sequence homologies may provide information about their evolutionary dynamics and potential functional conservation in related plant species. However, the level of conservation between previously published *B. rapa* lncRNAs and those identified in the present study was low, highlighting the lack of reported lncRNAs from male reproductive tissues in *B. napus* and tissue-specific expression of lncRNAs. In *A. thaliana*, Rutley and colleagues [49] reported novel pollen-specific lncRNAs and germinating pollen under controlled and heat-stressed conditions. Previous studies have also reported tissue or development-specific and species-specific expression of lncRNAs across various plants [48]. The expanding lncRNA database and comparative genomics may further the understanding of the functional conservation of lncRNAs, and the underlying mechanism across different tissues and plant species.

Recent investigations have revealed the various functions mediated by the action of lncRNAs in plants [50]. Different interactions between lncRNAs and protein-coding genes point toward the diverse mode of action of lncRNAs [33]. lncRNAs, like mRNA, can be miRNA targets and operate as miRNA decoys, suppressing the interaction between miRNAs and their target genes [13]. Out of the 1052 lincRNAs, only 22 were predicted as potential targets of 18 miRNAs, and 21 out of 780 lncNATs were predicted to be targeted by 36 miRNAs (Table S8). Some lncRNAs are also considered small RNA (miRNA and siRNA) precursors. We found only 10 lincRNA and 8 lncNATs are potential small RNA precursors (100% similarity to known mature miRNAs). The small fraction of lncRNAs identified as targets of miRNA and the lack of similarity between lncRNAs and mature miRNAs suggest that the majority of lncRNAs are unlikely to be miRNA decoys or act as small RNA precursors and have other, independent modes of regulation. However, it is notable that the publicly available databases may lack some miRNAs specific to pollen development; therefore, the roles of lncRNAs as miRNA decoys or miRNA precursors of yet unknown miRNAs cannot be excluded.

LncRNAs can regulate gene expression in a *cis*- or *trans*-manner [51]. Different classes of lncRNAs may play distinct roles in regulating protein-coding gene expression changes in relative abundance. lncRNAs act closer to the transcription site of the neighbouring genes while acting in a *cis* manner [52]. Contrary to this, they can regulate multiple genes throughout the genome by functioning away from the transcription site while acting in a *trans* manner [51]. The present study predicted *cis* and *trans* interactions between all the annotated lncRNAs and protein-coding genes (Figures 3 and 4, Table S2-S7). The *cis* interactions detected for lincRNAs and lncNATs highlighted that *cis*-acting lincRNAs are completely identified as intergenic. The distribution between sense/antisense and upstream/downstream description is approximately similar. *Cis*-acting lncNATs are mostly identified as antisense and genic, and most of them are located in exons of protein-coding genes. A higher number of significant *trans* interactions for lncNATs as compared to lincRNA were identified in this study. It is highly likely that the lncNATs overlapping with coding genes in the *B. rapa* genome potentially emerged from an ancestral genome triplication event [30, 53, 54]. A substantial proportion of *B. rapa* genes would be present in groups of paralogs having sequence similarity scattered in different positions on the genome [55]. Therefore, the chance of finding a high similarity between a lncNAT and a distant protein-coding gene is higher than for lincRNA, which is instead located outside protein-coding genes.

In plants, few studies have reported lncRNAs linked to male reproductive development, for instance, *LDMAR* in rice, *Zm401* in maize and *BcMF11* in *B. campestris [26-29]*. Huang et al. [42] also reported lncRNA-mRNA pairs, including 10 genes involved in pollen and pollen development functions. Weighted gene co-expression network and correlation analysis revealed 90 pairs of *cis*- or *trans*-acting lncRNAs and protein-coding genes, including several genes involved in transcriptional regulation, regulation of cell cycle, microtubule-based movement, pollen development, pollen tube growth, cell wall organization and transmembrane transport (Figure 6, 7 and Table S11). Interestingly, a higher number of *cis-*acting lncRNAs containing pairs were identified, and most of them were positively correlated (Table S11). The next obvious step is determining whether this link between *cis-*acting lncRNAs and protein-coding genes is causal. The intimate relationship between lncRNA/protein-coding gene pairs could indicate the presence of sophisticated gene expression regulatory mechanisms during pollen development. Several studies have reported *cis-*acting lncRNAs regulating the activation of neighbouring genes’ expression [21, 23, 56-59]. In our study, the lncRNAs that are strongly co-expressed with neighbouring protein-coding genes are thus promising candidates for further research. Even though cis-acting lncRNA and neighbouring protein-coding genes are typically positively correlated, we also found few lncRNA that potentially negatively regulated the expression of neighbouring genes. Several trans-acting linked lncRNAs were also discovered during pollen development. These results indicate that an intimate relationship between lncRNAs and protein-coding transcripts may be mediated by various molecular interactions.

## Material and Methods

### Plant material, sample collection and sequencing

*Brassica rapa* accession no. ATC 92270 Y.S (AND)-168 was used in this study. The plants were grown in a growth cabinet under the following conditions (21/18°C Day/night, a photoperiod of 16/8 hours light/dark, 200μmol m^-2^s^-1^ light intensity and 60% humidity). The different stages of pollen development were harvested as per previously published protocol [60, 61]. Briefly, the following five groups of buds were identified: <0.5mm buds (pollen/microspore mother cells, PMC), 0.8-1mm buds (tetrads, TET), 1-2.5mm (microspores to polarised microspores, MIC), 3-4.5mm (early to late bi-nucleate pollen, BIN) and 5-6mm (tri-nucleate pollen, POL). Anthers were carefully dissected from the buds of the last two groups and crushed in the B5 medium in a 1.5 mL tube. The whole buds were crushed in B5 medium for the first three groups. The crushed suspension was then filtered through a 44μm nylon mesh into 15 mL tubes. The filtrate was centrifuged at 150g for 3 min at 4°C. The supernatant was discarded, and the pellet was washed using 0.5 X B5 medium and was again centrifuged at 150g for 3 min at 4°C. The supernatant was removed, and the pellet was immediately frozen in liquid nitrogen and stored at -80°C. An aliquot from each isolation was analysed to check the developmental stage.

The buds collected from different plants were used as one biological replicate, and three independent biological replicates were prepared for each sample. The total RNA was isolated using the mirVana™ miRNA Isolation Kit (Thermo-Fisher; Part Numbers AM1560, AM1561, Carlsbad, CA, USA) according to the manufacturer’s instructions. The isolated RNA samples were treated with TURBO™DNase (Ambion, Carlsbad, CA, USA) to remove DNA contamination. The libraries were prepared using Illumina TruSeq stranded mRNA kit with poly (A) selection [61]. Additionally, five libraries were prepared using rRNA depletion [60]. The sequencing was performed at the Australian Genome Research Facility (AGRF), Melbourne.

### *B. rapa* genome re-annotation and lncRNA discovery

The reads were mapped to the *B. rapa* ‘Chiifu’ genome assembly v3.5 [30] using STARv2.7.9a [62] two-pass mapping as described by [63]. Default settings were used for generating the first-pass genome index. The filters and parameters for the two-pass mapping are provided in Table S1B. The transcripts for each library were assembled separately using Stringtie v2.1.4 [64, 65] using existing the *B. rapa* ‘Chiifu’ genome assembly v3.5 genome annotation as a guide. The individual assemblies were merged using Stringtie v2.1.4, again using the *B. rapa* ‘Chiifu’ genome assembly v3.5 genome annotation as a guide.

The coding potential of all genes was evaluated by (1) Coding potential calculator2, CPC2-beta, [66] (2) PLEK tool [67] (3) DIAMOND blastx v0.9.30 comparison against the RefSeq database (obtained on: 23.05.2021) [68] and (3) DIAMOND blastp comparison of the extracted longest open reading frames (ORFs; TransDecoder v5.5.0) against RefSeq database. The genes for which none of the transcripts was identified as coding were designated as non-coding. Non-coding loci with at least one transcript >=200 bp in length and no matched in the Rfam v14.7 database were identified as lncRNAs. The overlap between lncRNAs and protein-coding loci was identified by bedtools v2.30.0 [69]. lncRNAs that did not overlap any coding loci were designated lincRNAs, while lncRNA that overlapped at least one protein-coding locus was designated as lncNATs. A schematic representation of the pipeline for *B. rapa* genome re-annotation, lncRNA discovery and alternative splicing analysis is provided in Figure S1.

### *cis* and *trans* regulation of protein-coding genes by lncRNAs

Classification of lncRNAs acting in *cis* is performed with the FEELnc_classification script from the software FEELnc [70]. This software applies a 100kb sliding window and classifies lncRNAs based on its relationship with the closest mRNA in the window. A custom python script was used for generating statistics per chromosome of the annotated lncRNAs. The results are visualized in a Sankey plot with a custom R script using the packages ggplot2 and ggalluvial [71]. For simplicity, only the best-predicted interactions for lncRNAs localized in chromosomes are visualized, and the lncRNAs located on scaffold chromosomes were removed.

### Gene expression quantification, data pre-processing and differential expression analysis

Transcript expression was quantified using Kallisto v0.45.1 [72], and the downstream analysis was performed using the limma R package. Tximport R package version 1.10.0 (lengthScaledTPM method) was used to generate read counts and transcript per million reads (TPMs) [73]. Transcripts and genes with low expression were filtered based on data mean-variance trend analysis. A gene was considered expressed if any transcripts had CPM ≥ 1 in at least three of the 15 samples. Normalization of the gene and transcript read counts to *llllgg*_2_CPM was performed by using the TMM method [74]. Principal component analysis (PCA) was undertaken to determine the relatedness between the biological replicates (Figure S2A). Batch effects were corrected by using the RUVSeq R package [75]. Four contrast groups were defined to perform the differential expression analysis: TET vs PMC, MIC vs TET, BIN vs MIC and POL vs BIN. For a gene to be considered significantly differentially expressed in a contrast group, a cut-off adjusted p-value < 0.01 and log^2^ fold change ≥ 0.5 was used. P-values of multiple testing were adjusted to correct the false discovery rate (FDR) using the BH method [76].

### Functional annotation and enrichment analysis

Homologous *B. rapa* coding genes, compared to the Arabidopsis proteome, were identified using the BlastP program with an e-value ≤ 1e-05. PANNZER2 [77] was employed for the functional annotation of the protein-coding genes. PANNZER provides a GO term and a functional description for the query protein sequences. The GO enrichment analysis was performed using the GOstats R package, and a hypergeometric test (p-value < 0.01) was performed to identify overrepresented GO terms [78]. The annotated expressed genes were used as background for GO enrichment analysis. Further, ReViGO was used to retain the significant non-redundant GO terms [79].

Possible RNA-RNA interactions in *trans* between *B. rapa* lncRNAs and mRNAs were performed using the RlBLAST software [80]. RlBLAST was chosen for this analysis as it is over 64 times faster than other similar software for achieving a similar level of precision.

### lncRNA conservation analysis

Conservation analysis of lncRNAs, the sequences of identified lncRNA in this study were searched against lncRNAs downloaded from CANTATAdb 2.0 [32] for *A. thaliana, B. rapa, B. oleracea, B. napus, Glycine max, Solanum lycopersicum* and *Zea mays* with BLASTn [81]. The cut-off threshold of e-value 1e-10 and identity > 75% were employed to report the significant hits.

### Analysis of lncRNA as miRNA targets or precursors

The identification of *B. rapa* miRNA targets were done using psRNATarget v2 (2017 update) (Dai et al., 2018). Using the default parameters, the alignment of identified *B. rapa* lncRNAs and protein-coding genes expressed during pollen development was performed against the *B. rapa* miRNAs available on the psRNATarget database. A strict expectation threshold of ≤ 3 was used to filter potential targets.

Mature *B. rapa* sequences were downloaded from miRbase release 22.1 [82]. To identify whether lncRNAs acted as potential precursors or targets of small RNAs (miRNA and siRNA), the lncRNAs were compared to the miRNA sequences using BLASTN v2.7 (-task blastn - outfmt ‘6 qseqid sseqid pident length mismatch gapopen qstart qend sstart send evalue bitscore qcovs qlen slen’). The matches were filtered to retain only matches with 100% sequence identify and match length equal to miRNA. The matches were filtered to retain only matches with 100% sequence identify and match length equal to miRNA length.

### lncRNA-mRNA co-expression analysis by WGCNA

Weighted gene co-expression network analysis using the WGCNA R package v1.63 [83] was performed to identify coding genes and lncRNA co-expressed networks. As per the WGCNA guidelines, normalised gene counts were used. The construction of a co-expression network with an approximate scale-free topology of above 0.80 was achieved with a soft power of 10. A minimum module size of 30 is chosen. Moreover, gene modules that exhibit highly similar expression patterns are merged. After evaluating the module dendrogram, an arbitrary threshold of 0.20 (corresponding to 0.80 correlation) was set as limit to merge modules with a similar expression. The top three best correlating modules for each developmental stage were selected, and then the transcripts with the best significance in the selected correlated modules were identified, and functional enrichment analysis was performed. LncRNA-mRNA co-expressed pairs were identified among the hub genes of the selected modules and the potential regulatory relationship (cis or trans) was characterised between the pair. Further, the Pearson correlation coefficient between the lncRNA-protein coding gene co-expressed pairs was calculated using the R package.

## Conclusions

In summary, we investigated the expression of long non-coding RNAs during pollen development and identified 1,832 lncRNA (1052 lincRNA, 780 lncNAT) and 31,729 protein-coding genes as expressed during pollen development in *B. rapa*. The lncRNAs have defined stage-specific expression patterns. The lncRNAs were subdivided into classes based on their genomic location and orientation relative to protein-coding gene neighbours. lncRNA belonging to different classes have distinct properties suggesting possible differences in function and/or mode of action. The analysis of expression patterns of lncRNA-protein coding genes points to the involvement of lncRNAs in the modulation of the protein-coding gene associated with several biological processes regulating pollen development. Overall, genome-wide identification, characterisation and functional analysis enabled the identification of lncRNAs candidates and their functional associations with protein-coding genes, potentially revealing regulatory and molecular mechanisms underlying male reproductive development in *B. rapa*.

## Author Contributions

Samples collection and RNA isolation, A.D.A; data curation and analysis, N.L. and A.A.G.; initial draft preparation, N.L. and A.A.G.; review and editing, M.B.S. and P.L.B.; supervision, M.B.S. and P.L.B. All authors have read and agreed to the published version of the manuscript.

## Institutional Review Board Statement

Not applicable.

## Informed Consent Statement

Not applicable.

## Data Availability Statement

All data generated in this study are available in the article and its Supplementary Materials. The Poly-A strand specific RNA-Sequencing data were deposited in NCBI SRA under PRJNA529957 and rRNA depletion RNA-Seq data were deposited in NCBI SRA under PRJNA763698.

## Acknowledgments

This research was supported by Spartan HPC (Lev Lafayette GS, Linh Vu, Bernard Meade October 27, 2016 [84]) at the University of Melbourne, Australia.

## References

1. Loraine, A.E., et al., RNA-seq of Arabidopsis pollen uncovers novel transcription and alternative splicing. Plant physiology, 2013. 162(2): p. 1092–1109.

2. Honys, D. and D. Twell, Transcriptome analysis of haploid male gametophyte development in Arabidopsis. Genome biology, 2004. 5(11): p. 1–13.

3. Rutley, N. and D. Twell, A decade of pollen transcriptomics. Plant reproduction, 2015. 28(2): p. 73–89.

4. Suwabe, K., et al., Separated transcriptomes of male gametophyte and tapetum in rice: validity of a laser microdissection (LM) microarray. Plant and cell physiology, 2008. 49(10): p. 1407–1416.

5. Borg, M., L. Brownfield, and D. Twell, Male gametophyte development: a molecular perspective. Journal of experimental botany, 2009. 60(5): p. 1465–1478.

6. Twell, D., S.-A. Oh, and D. Honys, Pollen development, a genetic and transcriptomic view, in The pollen tube. 2006, Springer. p. 15–45.

7. da Costa-Nunes, J.A. and U. Grossniklaus, Unveiling the gene-expression profile of pollen. Genome biology, 2003. 5(1): p. 1–3.

8. Brownfield, L., et al., A plant germline-specific integrator of sperm specification and cell cycle progression. PLoS genetics, 2009. 5(3): p. e1000430.

9. Perry, R.B.-T. and I. Ulitsky, The functions of long noncoding RNAs in development and stem cells. Development, 2016. 143(21): p. 3882–3894.

10. Golicz, A.A., P.L. Bhalla, and M.B. Singh, lncRNAs in plant and animal sexual reproduction. Trends in plant science, 2018. 23(3): p. 195–205.

11. Mattick, J.S. and J.L. Rinn, Discovery and annotation of long noncoding RNAs. Nature structural & molecular biology, 2015. 22(1): p. 5–7.

12. Wu, H.-J., et al., Widespread long noncoding RNAs as endogenous target mimics for microRNAs in plants. Plant physiology, 2013. 161(4): p. 1875–1884.

13. Franco-Zorrilla, J.M., et al., Target mimicry provides a new mechanism for regulation of microRNA activity. Nature genetics, 2007. 39(8): p. 1033–1037.

14. Wang, K.C. and H.Y. Chang, Molecular mechanisms of long noncoding RNAs. Molecular cell, 2011. 43(6): p. 904–914.

15. Böhmdorfer, G. and A.T. Wierzbicki, Control of chromatin structure by long noncoding RNA. Trends in cell biology, 2015. 25(10): p. 623–632.

16. Zhang, Y.-C., et al., Genome-wide screening and functional analysis identify a large number of long noncoding RNAs involved in the sexual reproduction of rice. Genome biology, 2014. 15(12): p. 1–16.

17. Li, S., et al., High-resolution expression map of the Arabidopsis root reveals alternative splicing and lincRNA regulation. Developmental cell, 2016. 39(4): p. 508–522.

18. Di, C., et al., Characterization of stress-responsive lnc RNA s in A rabidopsis thaliana by integrating expression, epigenetic and structural features. The Plant Journal, 2014. 80(5): p. 848–861.

19. Yuan, J., et al., Systematic characterization of novel lncRNAs responding to phosphate starvation in Arabidopsis thaliana. BMC genomics, 2016. 17(1): p. 1–16.

20. Yu, X., et al., Global analysis of cis-natural antisense transcripts and their heat-responsive nat-siRNAs in Brassica rapa. BMC Plant Biology, 2013. 13(1): p. 1–13.

21. Csorba, T., et al., Antisense COOLAIR mediates the coordinated switching of chromatin states at FLC during vernalization. Proceedings of the National Academy of Sciences, 2014. 111(45): p. 16160–16165.

22. Heo, J.B. and S. Sung, Vernalization-mediated epigenetic silencing by a long intronic noncoding RNA. Science, 2011. 331(6013): p. 76–79.

23. Rosa, S., S. Duncan, and C. Dean, Mutually exclusive sense–antisense transcription at FLC facilitates environmentally induced gene repression. Nature Communications, 2016. 7(1): p. 1–7.

24. Zhao, X., et al., Global identification of Arabidopsis lncRNAs reveals the regulation of MAF4 by a natural antisense RNA. Nature communications, 2018. 9(1): p. 1–12.

25. Wang, Y., et al., Overexpressing lncRNA LAIR increases grain yield and regulates neighbouring gene cluster expression in rice. Nature communications, 2018. 9(1): p. 1–9.

26. Ding, J., et al., A long noncoding RNA regulates photoperiod-sensitive male sterility, an essential component of hybrid rice. Proceedings of the National Academy of Sciences, 2012. 109(7): p. 2654–2659.

27. Ma, J., et al., Zm401, a short-open reading-frame mRNA or noncoding RNA, is essential for tapetum and microspore development and can regulate the floret formation in maize. Journal of cellular biochemistry, 2008. 105(1): p. 136–146.

28. Song, J.-H., J.-S. Cao, and C.-G. Wang, BcMF11, a novel non-coding RNA gene from Brassica campestris, is required for pollen development and male fertility. Plant cell reports, 2013. 32(1): p. 21–30.

29. Song, J.-H., et al., BcMF11, a putative pollen-specific non-coding RNA from Brassica campestris ssp. chinensis. Journal of plant physiology, 2007. 164(8): p. 1097–1100.

30. Zhang, Z., et al., Improved Reference Genome Annotation of Brassica rapa by Pacific Biosciences RNA Sequencing. Frontiers in plant science, 2022. 13: p. 841618.

31. Golicz, A., Long Intergenic Noncoding RNA (lincRNA) Discovery from Non-Strand-Specific RNA-Seq Data, in Plant Bioinformatics. 2022, Springer. p. 465-482.

32. Szcześniak, M.W., et al., CANTATAdb 2.0: expanding the collection of plant long noncoding RNAs, in Plant Long Non-Coding RNAs. 2019, Springer. p. 415–429.

33. Rinn, J.L. and H.Y. Chang, Genome regulation by long noncoding RNAs. Annual review of biochemistry, 2012. 81: p. 145–166.

34. Kornienko, A.E., et al., Gene regulation by the act of long non-coding RNA transcription. BMC biology, 2013. 11(1): p. 1–14.

35. Bartel, D.P., MicroRNAs: genomics, biogenesis, mechanism, and function. cell, 2004. 116(2): p. 281–297.

36. Wei, L.Q., et al., Genome-scale analysis and comparison of gene expression profiles in developing and germinated pollen in Oryza sativa. Bmc Genomics, 2010. 11(1): p. 1–20.

37. Waseem, M., Y. Liu, and R. Xia, Long non-coding RNAs, the dark matter: an emerging regulatory component in plants. International Journal of Molecular Sciences, 2020. 22(1): p. 86.

38. Yu, Y., et al., Plant noncoding RNAs: hidden players in development and stress responses. Annual review of cell and developmental biology, 2019. 35: p. 407.

39. Ward, M., et al., Conservation and tissue-specific transcription patterns of long noncoding RNAs. Journal of human transcriptome, 2015. 1(1): p. 2–9.

40. Gawronski, K.A. and J. Kim, Single cell transcriptomics of noncoding RNAs and their cell-specificity. Wiley Interdisciplinary Reviews: RNA, 2017. 8(6): p. e1433.

41. Wang, M., et al., Long noncoding RNA s and their proposed functions in fibre development of cotton (Gossypium spp.). New phytologist, 2015. 207(4): p. 1181–1197.

42. Huang, L., et al., Systematic identification of long non-coding RNA s during pollen development and fertilization in Brassica rapa. The Plant Journal, 2018. 96(1): p. 203–222.

43. Golicz, A.A., M.B. Singh, and P.L. Bhalla, The long intergenic noncoding RNA (LincRNA) landscape of the soybean genome. Plant Physiology, 2018. 176(3): p. 2133–2147.

44. Liu, J., et al., Genome-wide analysis uncovers regulation of long intergenic noncoding RNAs in Arabidopsis. The Plant Cell, 2012. 24(11): p. 4333–4345.

45. Smith, M.A. and J.S. Mattick, Structural and functional annotation of long noncoding RNAs, in Bioinformatics. 2017, Springer. p. 65–85.

46. Barreau, C., L. Paillard, and H.B. Osborne, AU-rich elements and associated factors: are there unifying principles? Nucleic acids research, 2005. 33(22): p. 7138–7150.

47. Simopoulos, C.M., E.A. Weretilnyk, and G.B. Golding, Molecular traits of long non-protein coding RNAs from diverse plant species show little evidence of phylogenetic relationships. G3: Genes, Genomes, Genetics, 2019. 9(8): p. 2511–2520.

48. Ke, L., et al., Evolutionary dynamics of linc RNA transcription in nine citrus species. The Plant Journal, 2019. 98(5): p. 912–927.

49. Rutley, N., et al., Characterization of novel pollen-expressed transcripts reveals their potential roles in pollen heat stress response in Arabidopsis thaliana. Plant reproduction, 2021. 34(1): p. 61–78.

50. Nejat, N. and N. Mantri, Emerging roles of long non-coding RNAs in plant response to biotic and abiotic stresses. Critical reviews in biotechnology, 2018. 38(1): p. 93–105.

51. Fatica, A. and I. Bozzoni, Long non-coding RNAs: new players in cell differentiation and development. Nature Reviews Genetics, 2014. 15(1): p. 7–21.

52. Guil, S. and M. Esteller, Cis-acting noncoding RNAs: friends and foes. Nature structural & molecular biology, 2012. 19(11): p. 1068.

53. Cheng, F., J. Wu, and X. Wang, Genome triplication drove the diversification of Brassica plants. Horticulture research, 2014. 1.

54. Cai, C., et al., Brassica rapa genome 2.0: a reference upgrade through sequence re-assembly and gene re-annotation. Molecular plant, 2017. 10(4): p. 649–651.

55. Mun, J.-H., et al., Genome-wide comparative analysis of the Brassica rapa gene space reveals genome shrinkage and differential loss of duplicated genes after whole genome triplication. Genome biology, 2009. 10(10): p. 1–18.

56. Vance, K.W. and C.P. Ponting, Transcriptional regulatory functions of nuclear long noncoding RNAs. Trends in Genetics, 2014. 30(8): p. 348–355.

57. Statello, L., et al., Gene regulation by long non-coding RNAs and its biological functions. Nature reviews Molecular cell biology, 2021. 22(2): p. 96–118.

58. Krishnan, J. and R.K. Mishra, Emerging trends of long non-coding RNA s in gene activation. The FEBS journal, 2014. 281(1): p. 34–45.

59. Yap, K.L., et al., Molecular interplay of the noncoding RNA ANRIL and methylated histone H3 lysine 27 by polycomb CBX7 in transcriptional silencing of INK4a. Molecular cell, 2010. 38(5): p. 662–674.

60. Babaei, S., M.B. Singh, and P.L. Bhalla, Circular RNAs repertoire and expression profile during Brassica rapa pollen development. International journal of molecular sciences, 2021. 22(19): p. 10297.

61. Golicz, A.A., et al., A dynamic intron retention program regulates the expression of several hundred genes during pollen meiosis. Plant Reproduction, 2021. 34(3): p. 225–242.

62. Dobin, A. and T.R. Gingeras, Mapping RNA-seq reads with STAR. Current protocols in bioinformatics, 2015. 51(1): p. 11.14. 1-11.14. 19.

63. Veeneman, B.A., et al., Two-pass alignment improves novel splice junction quantification. Bioinformatics, 2016. 32(1): p. 43–49.

64. Pertea, M., et al., StringTie enables improved reconstruction of a transcriptome from RNA-seq reads. Nature biotechnology, 2015. 33(3): p. 290–295.

65. Varabyou, A., S.L. Salzberg, and M. Pertea, Effects of transcriptional noise on estimates of gene and transcript expression in RNA sequencing experiments. Genome research, 2021. 31(2): p. 301–308.

66. Kang, Y.-J., et al., CPC2: a fast and accurate coding potential calculator based on sequence intrinsic features. Nucleic acids research, 2017. 45(W1): p. W12–W16.

67. Li, A., J. Zhang, and Z. Zhou, PLEK: a tool for predicting long non-coding RNAs and messenger RNAs based on an improved k-mer scheme. BMC bioinformatics, 2014. 15(1): p. 1–10.

68. Buchfink, B., C. Xie, and D.H. Huson, Fast and sensitive protein alignment using DIAMOND. Nature methods, 2015. 12(1): p. 59–60.

69. Quinlan, A.R., BEDTools: the Swiss-army tool for genome feature analysis. Current protocols in bioinformatics, 2014. 47(1): p. 11.12. 1-11.12. 34.

70. Wucher, V., et al., FEELnc: a tool for long non-coding RNA annotation and its application to the dog transcriptome. Nucleic acids research, 2017. 45(8): p. e57–e57.

71. Wickham, H., ggplot2: elegant graphics for data analysis. 2016: springer.

72. Bray, N.L., et al., Near-optimal probabilistic RNA-seq quantification. Nature biotechnology, 2016. 34(5): p. 525–527.

73. Soneson, C., M.I. Love, and M.D. Robinson, Differential analyses for RNA-seq: transcript-level estimates improve gene-level inferences. F1000Research, 2015. 4.

74. Bullard, J.H., et al., Evaluation of statistical methods for normalization and differential expression in mRNA-Seq experiments. BMC bioinformatics, 2010. 11(1): p. 1–13.

75. Risso, D., RUVSeq: remove unwanted variation from RNA-seq data. Bioconductor https://bioconductor.org/packages/release/bioc/html/RUVSeq.html, 2015.

76. Benjamini, Y. and D. Yekutieli, The control of the false discovery rate in multiple testing under dependency. Annals of statistics, 2001: p. 1165–1188.

77. Törönen, P., A. Medlar, and L. Holm, PANNZER2: a rapid functional annotation web server. Nucleic acids research, 2018. 46(W1): p. W84–W88.

78. Falcon, S. and R. Gentleman, Using GOstats to test gene lists for GO term association. Bioinformatics, 2007. 23(2): p. 257–258.

79. Supek, F., et al., REVIGO summarizes and visualizes long lists of gene ontology terms. PloS one, 2011. 6(7): p. e21800.

80. Fukunaga, T. and M. Hamada, RIblast: an ultrafast RNA–RNA interaction prediction system based on a seed-and-extension approach. Bioinformatics, 2017. 33(17): p. 2666–2674.

81. Chen, Y., et al., High speed BLASTN: an accelerated MegaBLAST search tool. Nucleic acids research, 2015. 43(16): p. 7762–7768.

82. Kozomara, A., M. Birgaoanu, and S. Griffiths-Jones, miRBase: from microRNA sequences to function. Nucleic acids research, 2019. 47(D1): p. D155–D162.

83. Langfelder, P. and S. Horvath, WGCNA: an R package for weighted correlation network analysis. BMC bioinformatics, 2008. 9(1): p. 1–13.

84. Meade, B., et al., Spartan HPC-cloud hybrid: delivering performance and flexibility. University of Melbourne, 2017. 10: p. 49.

