## Supplementary material for "Genome-wide analysis of lncRNAs points to their roles in the modulation of developmental regulator expression during plant male germline development": Figure S

**Figure S1** Pipeline for reference-based transcriptome assembly and the identification of lncRNAs in *Brassica rapa*.

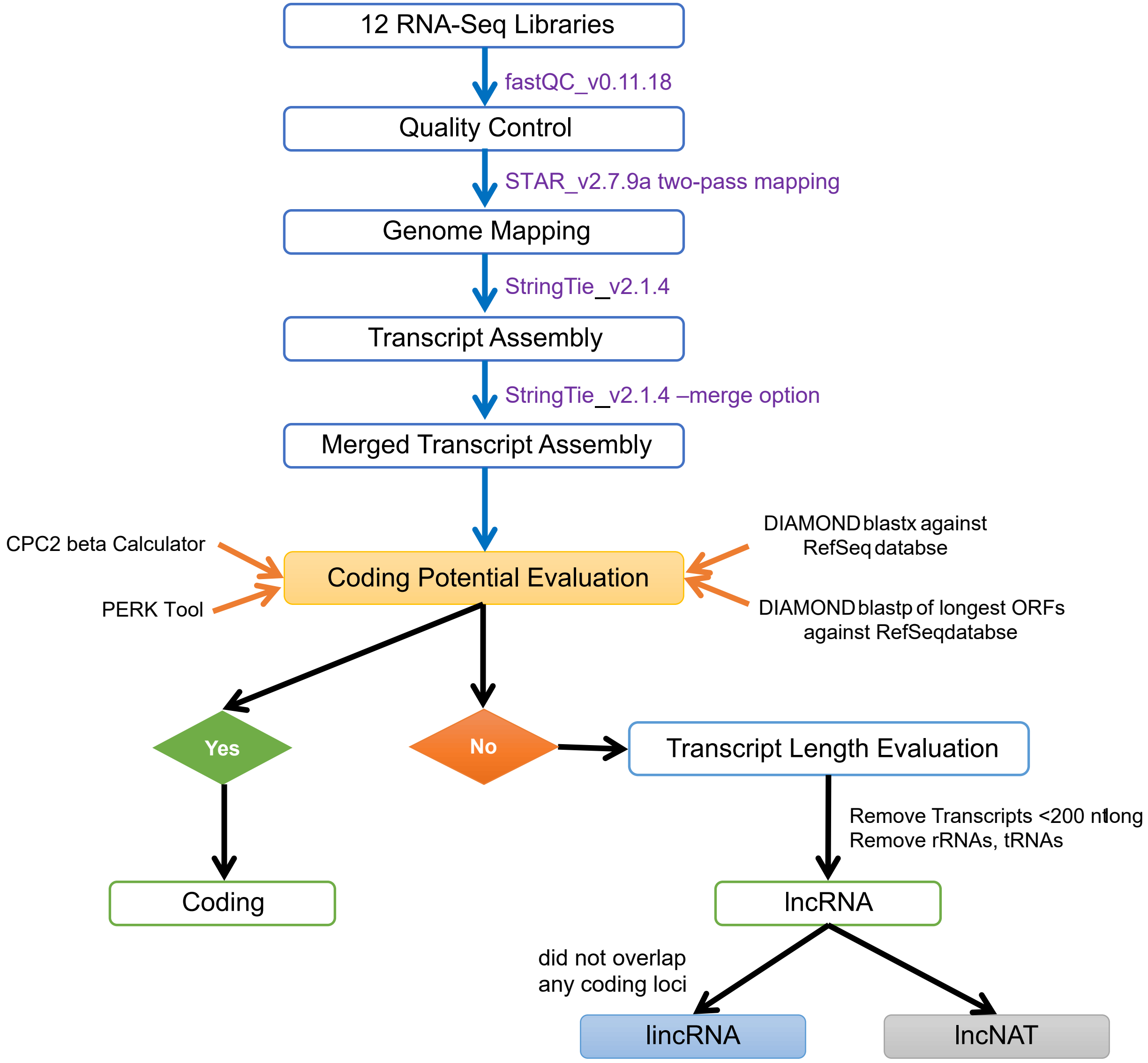

**Figure S2 (A)** Principal Component Analysis (PCA) between three biological replicates of each sample (batch effects removed),**(B)** Pearson correlation between expression of coding, lincRNA and lncNAT genes across the samples, and **(C)** Details and TPM values of annotated known pollen development marker genes. PMC: pollen/ microspore mother cell, TET: tetrads, MIC: microspores to polarised microspores, BIN: early to late bi-nucleate pollen, and POL: tri-nucleate pollen.

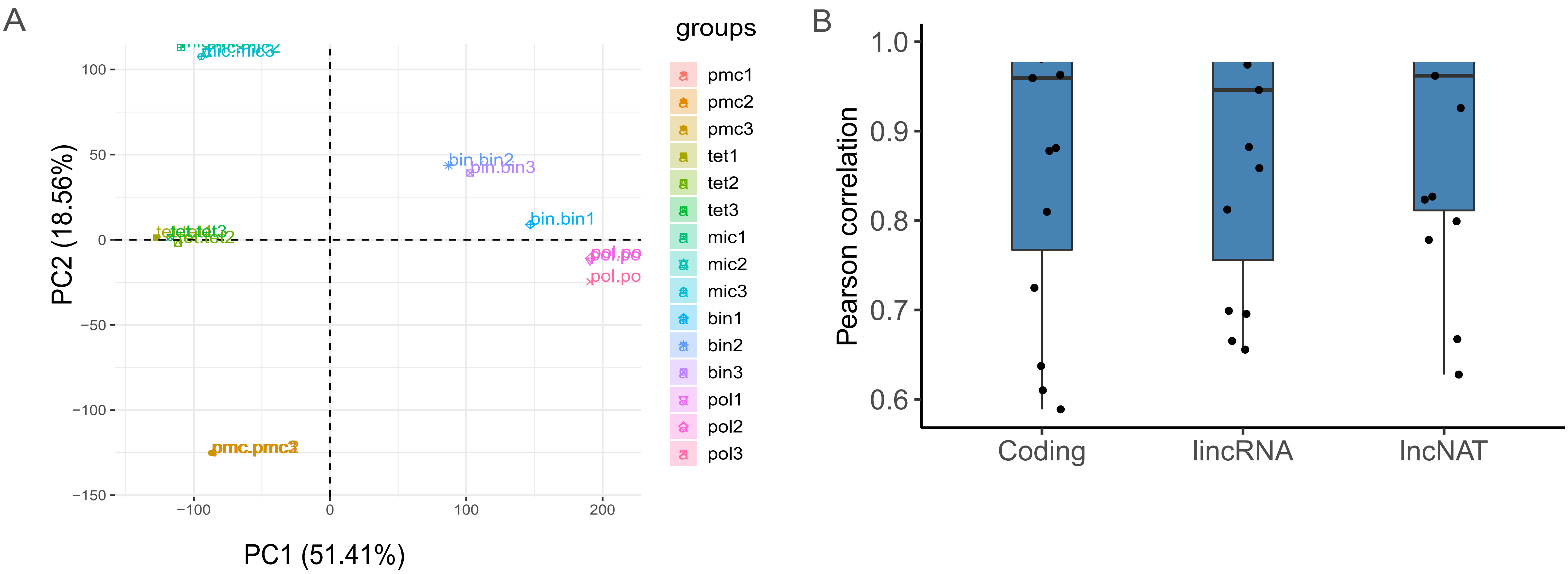


C

| Gene_name | *Arabidopsis* _ID | *B. rapa* ID | PMC | TET | MIC | BIN | POL |
|---|---|---|---|---|---|---|---|
| AKV | AT4G05440 | BRAPST00035771 | 24.76 | 23.32 | 30.42 | 35.12 | 14.92 |
| AtMGH3 | AT1G19890 | N/A | N/A | N/A | N/A | N/A | N/A |
| DUO1 | AT3G60460 | BRAPST00030023 | 0.06 | 0.33 | 0.45 | 29.26 | 15.97 |
| EMS1 | AT5G07280 | BRAPST00044981 | 22.76 | 10.64 | 5.18 | 0.15 | 0.02 |
| FBL17 | AT3G54650 | BRAPST00016105 | 5.09 | 4.08 | 3.05 | 1.80 | 0.22 |
| GEX1a | AT5G55490 | BRAPST00010468 | 0.19 | 1.52 | 1.67 | 1.45 | 6.88 |
| GEX1b | AT5G55490 | BRAPST00043379 | 0.93 | 11.26 | 13.37 | 0.59 | 9.09 |
| GEX3 | AT5G16020 | BRAPST00044352 | 0.22 | 0.20 | 0.03 | 6.27 | 10.92 |
| HAP2a | AT4G11720 | BRAPST00038234 | 0.18 | 2.02 | 1.06 | 16.71 | 23.20 |
| HAP2b | AT4G11720 | BRAPST00040134 | 32.39 | 14.90 | 7.91 | 22.79 | 9.57 |
| HAP2c | AT4G11720 | BRAPST00040601 | 0.18 | 0.64 | 0.51 | 73.55 | 25.71 |

**Figure S3 (A)** Number of up- and down-regulated gene in the four contrasts: TET-PMC, MIC-TET, BIN-MIC and POL-BIN, **(B)** Venn diagrams showing extent of overlap between the up- and down-regulated coding genes and lncRNAs across the four contrasts: TET-PMC, MIC-TET, BIN-MIC and POL-BIN (PMC: pollen/ microspore mother cell, TET: tetrads, MIC: microspores to polarised microspores, BIN: early to late bi-nucleate pollen, and POL: tri-nucleate pollen).

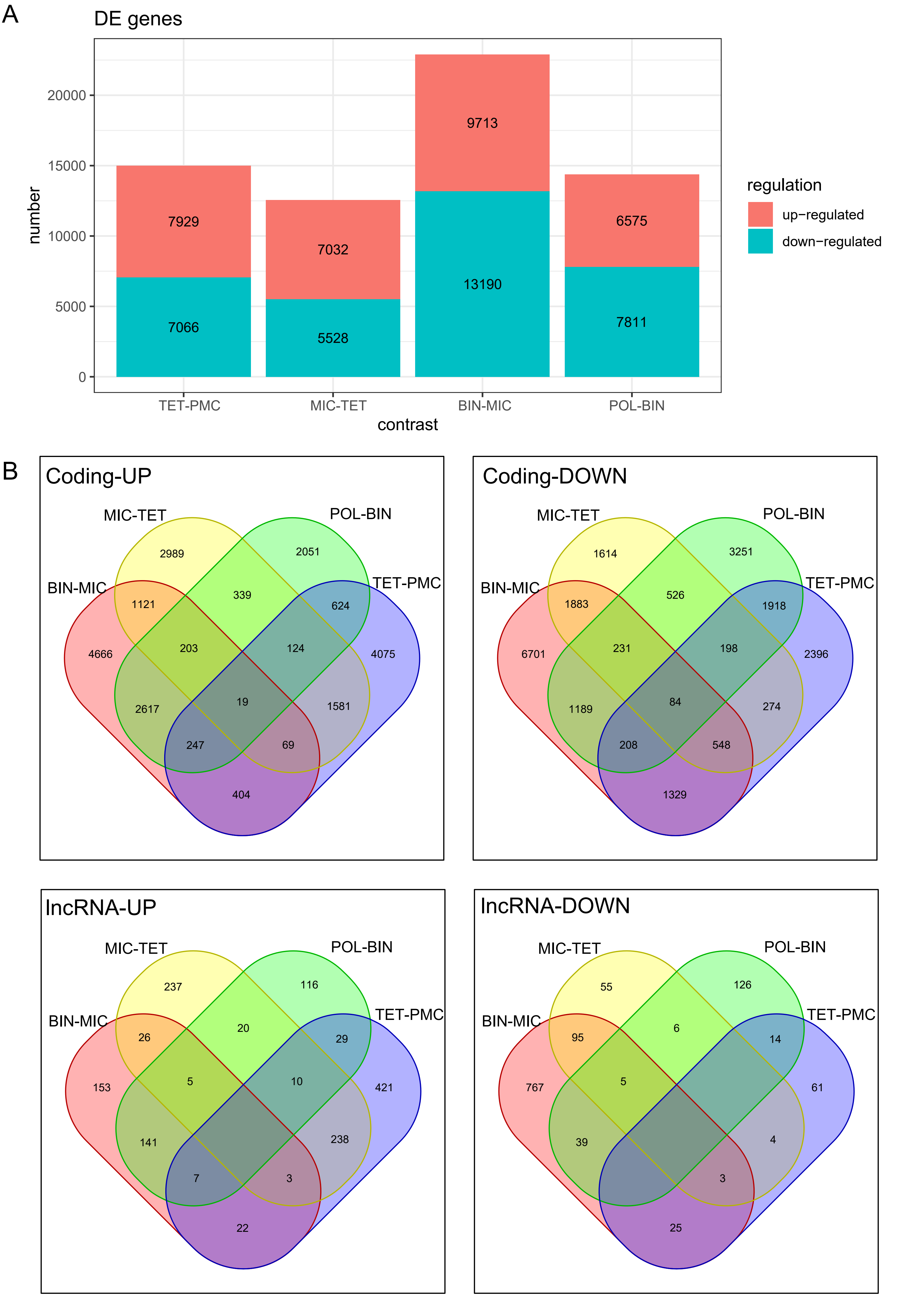

**Figure S4** GO enrichment of differentially regulated genes **(A)** Top 30 non-redundant GO biological processes terms associated with up-regulated genes across the four contrasts (TET-PMC, MIC-TET, BIN-MIC and POL-BIN), and **(B)** op 30 non-redundant GO biological processes terms associated with down-regulated genes across the four contrasts (TET-PMC, MIC-TET, BIN-MIC and POL-BIN), where PMC: pollen/ microspore mother cell, TET: tetrads, MIC: microspores to polarised microspores, BIN: early to late bi-nucleate pollen, and POL tri-nucleate pollen.

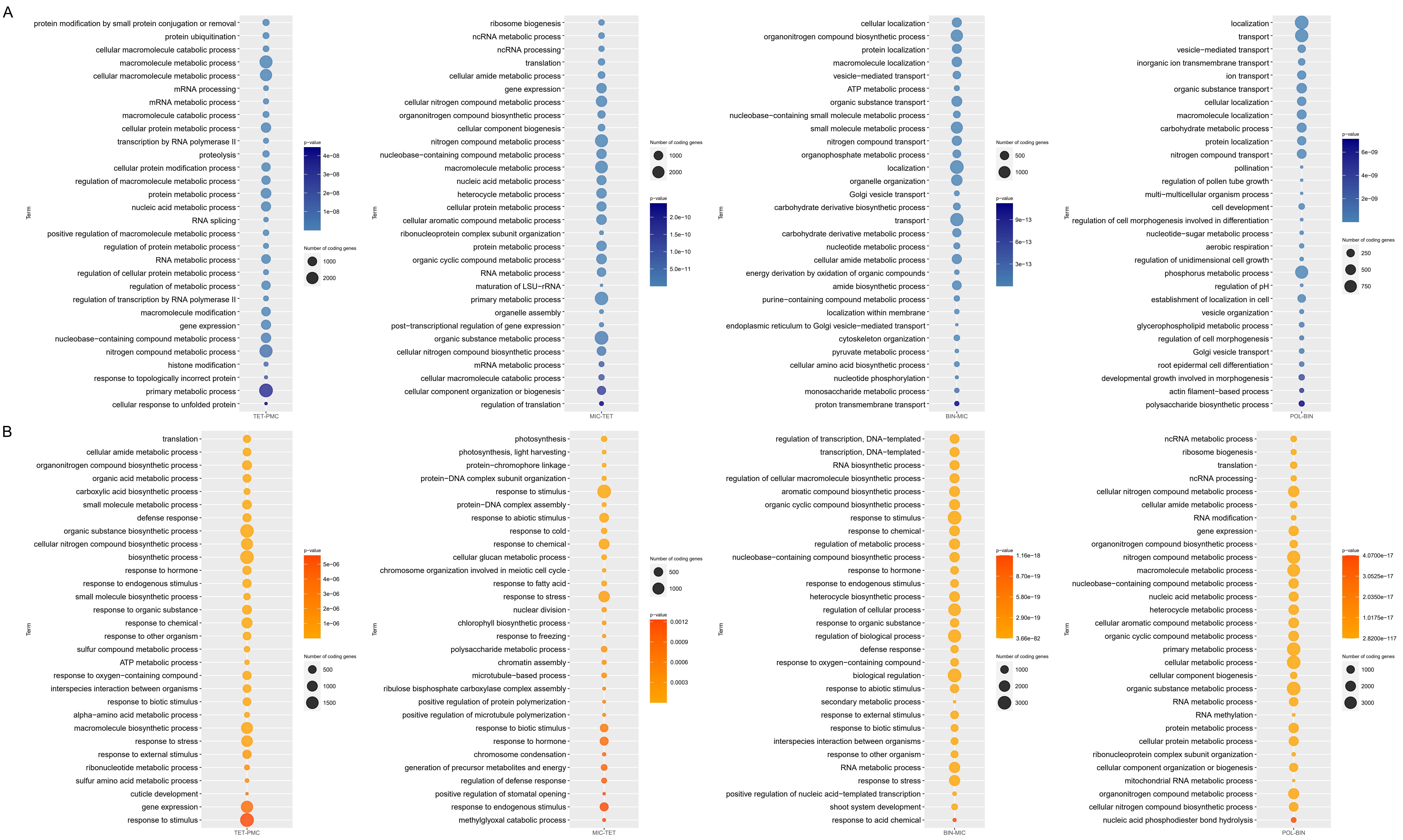

**Figure S5 (A)** Samples hierarchical clustering of $\log_2$ normalised gene counts and correlation with mock traits, **(B)** Comparison of dynamic tree cut and merged dynamic modules, and **(C)** Module-sample traits correlation heatmap.

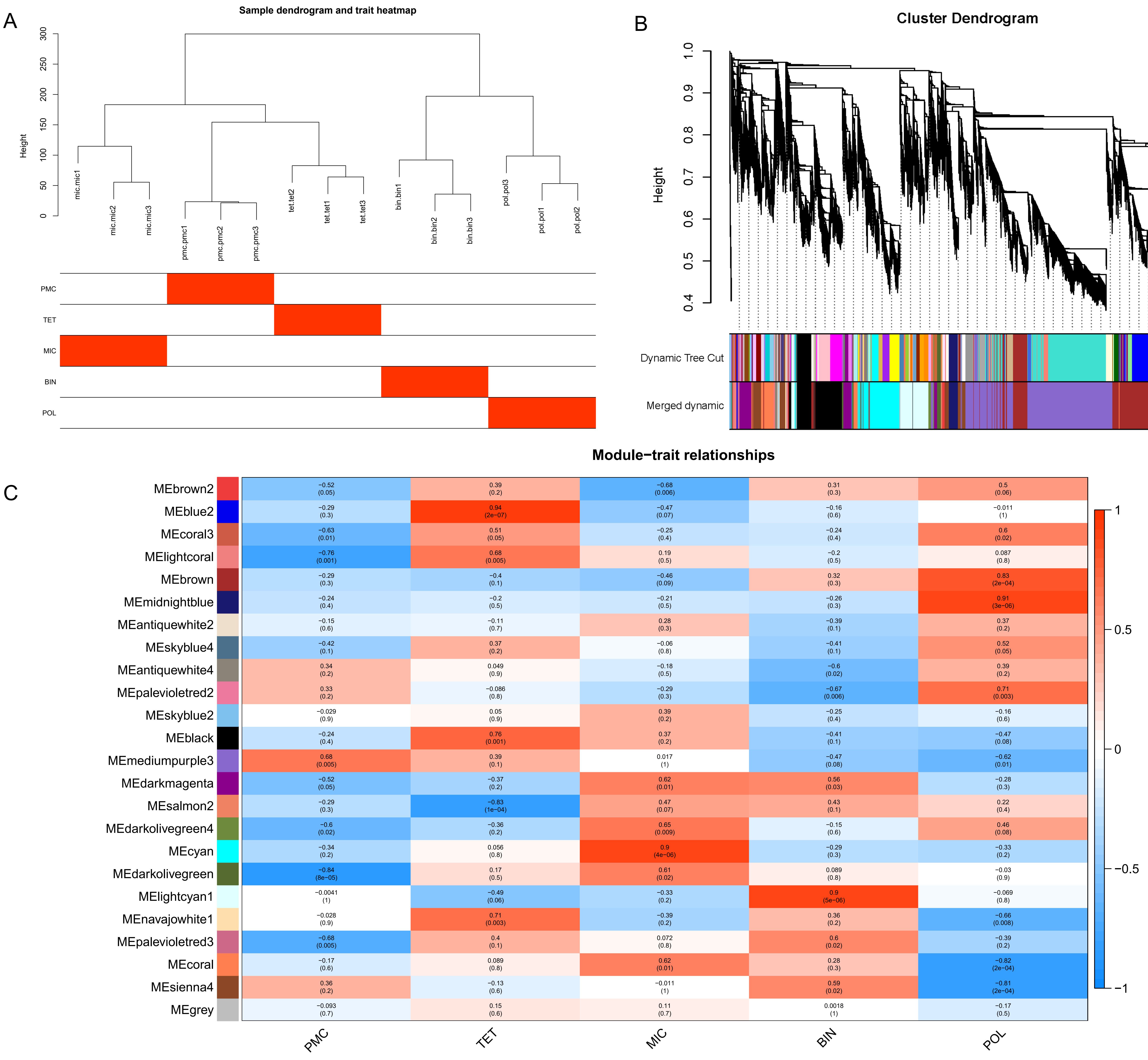
